# Auxin-driven ecophysiological diversification of leaves in domesticated tomato

**DOI:** 10.1101/2021.12.16.473023

**Authors:** Juliene d. R. Moreira, Bruno L. Rosa, Bruno S. Lira, Joni E. Lima, Ludmila N. Souza, Wagner C. Otoni, Antonio Figueira, Luciano Freschi, Tetsu Sakamoto, Lázaro E. P. Peres, Magdalena Rossi, Agustin Zsögön

**Author notes:** **For correspondence**: Agustin Zsögön. **Author contributions:** JEL, LEPP, MR, AZ conceived the project, JDRM, BLR, BSL, JEL, LNS, TS performed experiments; LF, WCO, AF, LEPP, MR, AZ supervised experiments; JDRM and AZ wrote the article with contributions from all the authors; AZ agrees to serve as the author responsible for contact. **Funding:** Fundação de Amparo à Pesquisa do Estado de São Paulo (FAPESP), Brazil 2016/01128-9; Fundação de Amparo à Pesquisa do Estado de São Paulo (FAPESP), Brazil 2017/14953-0; Conselho Nacional de Desenvolvimento Científico e Tecnológico (CNPq); Coordenação de Aperfeiçoamento de Pessoal de Nível Superior (CAPES), Brazil, Finance Code 001; Foundation for Research Assistance of the Minas Gerais State (FAPEMIG), Brazil RED-00053-16.

## Abstract

Heterobaric leaves have bundle sheath extensions (BSEs) that compartmentalise the parenchyma, whereas homobaric leaves do not. The presence of BSEs affects leaf hydraulics and photosynthetic rate. The tomato (*Solanum lycopersicum*) *obscuravenosa* (*obv*) mutant lacks BSEs. Here we identify the *obv* gene and the causative mutation, a non-synonymous amino acid change that disrupts a C2H2 zinc finger motif in a putative transcription factor. This mutation exists as a rare polymorphism in the natural range of wild tomatoes, but has increased in frequency in domesticated tomatoes, suggesting that the latter diversified into heterobaric and homobaric leaf types. The *obv* mutant displays reduced vein density, leaf hydraulic conductance and photosynthetic assimilation rate. We show that these and other effects on plant development, including changes in leaf insertion angle, leaf margin serration, minor vein density and fruit shape, are controlled by OBV via changes in auxin signalling. Loss of function of the transcriptional regulator AUXIN RESPONSE FACTOR (ARF4) also results in defective BSE development, revealing an additional component of a novel genetic module controlling aspects of leaf development important for ecological adaptation and subject to breeding selection.

**One sentence summary:** distribution of tomato heterobaric and homobaric leaves is controlled by a single-nucleotide polymorphism in an auxin-related transcription factor

## INTRODUCTION

Crop domestication was driven by artificial selection to create a loose suite of traits known as the ‘domestication syndrome’ (Hammer, 1984). Considerable effort has been devoted to unveiling the genetic basis of these traits. However, the passage from natural ecosystems to human-managed ones created new drivers for crop evolution that may have had impact beyond the domestication syndrome (Milla et al., 2015). These drivers include geographic expansion beyond the crops’ centre of origin, natural selection under cultivation in a highly managed environment, and indirect selection due to constraints caused by developmental trait correlations and physiological trade-offs (Milla et al., 2014). The extent and breadth of the genetic signature created by these processes in crops is not yet known. Understanding the genetic and physiological basis of functionally important traits would provide insights for both agricultural and evolutionary studies (Moyle and Muir, 2010). It would further provide valuable support to crop breeding and to the effort of creating new crops using *de novo* domestication pipelines (Zsögön et al., 2018; Gasparini et al., 2021).

Leaf functional traits are key for resource acquisition and are thus strongly constrained by developmental co-variation (Wright et al., 2004). For instance, high photosynthetic rates requires high transpiration capacity, which in turn depends on water transport capacity (Brodribb et al., 2007). Leaf hydraulic conductance is a fundamental driver of leaf diversity, as it is linked with leaf shape, longevity and venation architecture (Sack and Holbrook, 2006). Notably, many angiosperm species have translucent leaf veins due to the presence of bundle sheath extensions (BSEs), compact columns of chlorophyll-less cells that link the veins to the leaf epidermis (Haberlandt, 1882; Wylie, 1952) (Figure 1A). BSEs have considerable functional impact, as they can divide the leaf mesophyll into compartments, resulting in patchy stomatal opening and non-uniform photosynthesis (Terashima, 1992). Leaves harboring such compartments created by BSEs are called heterobaric, as opposed to homobaric ones, where the leaf lamina is topologically continuous (Pieruschka et al., 2010). The effect of BSEs on leaf compartmentalization can be demonstrated by water infiltration in the lamina (Kenzo et al., 2007).

**Figure 1.**
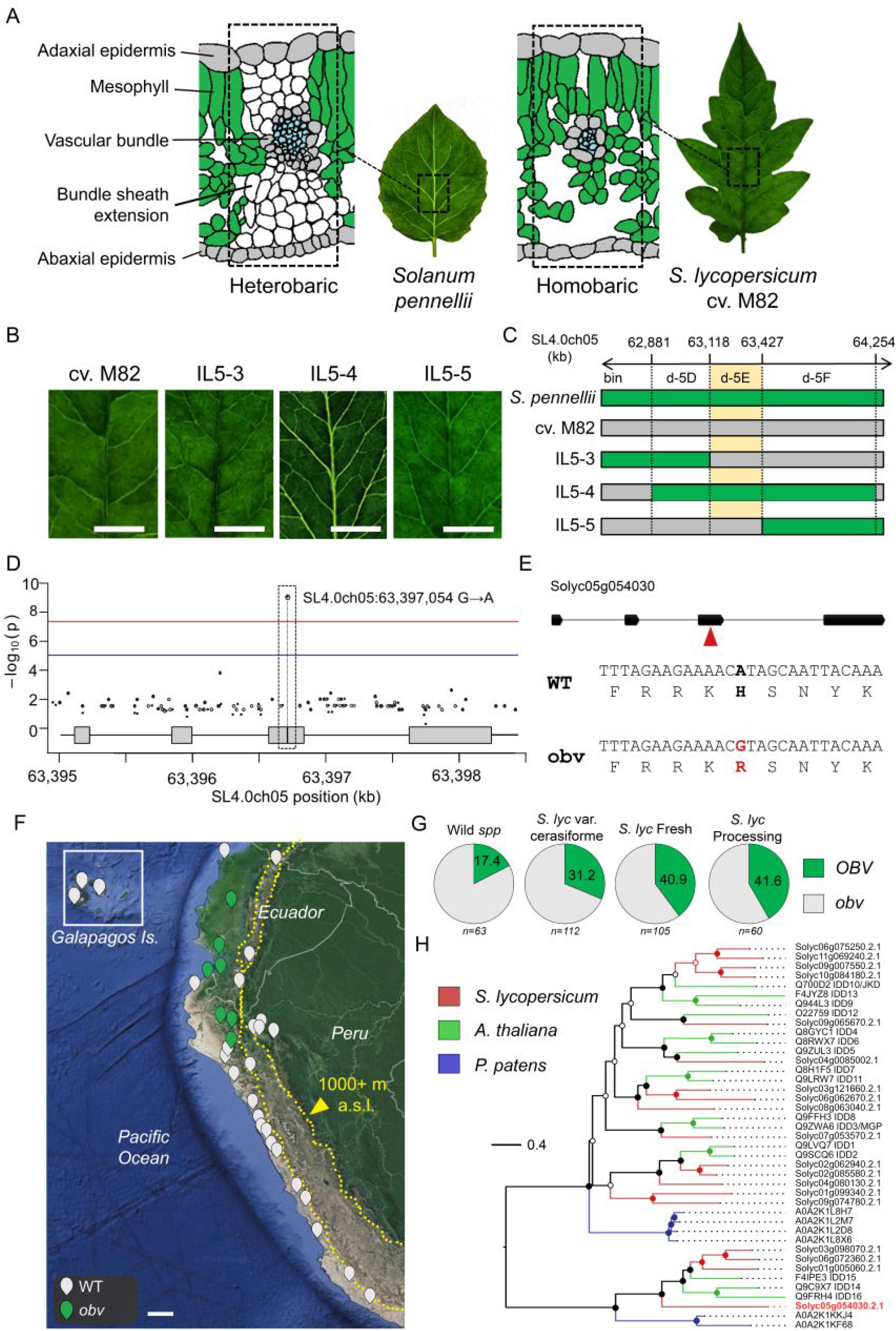
Mapping and identification of a candidate gene for the *obscuravenosa* (*obv*) mutation. (**A**) Schematic cross-sectional comparison of heterobaric and homobaric leaves. In the former (*e*.*g*. the tomato wild relative *Solanum pennellii*) the bundle sheath extensions (BSEs) are visible as a translucent network of veins, whereas leaves of the latter (*e*.*g*. cultivars of tomato, *S. lycopersicum*, harboring the *obv* mutation, such as cv. M82) the leaf lamina appears uniformly green. (**B**) Leaf phenotype of tomato cv. M82 and the *S. pennellii* introgression lines 5-3, 5-4 and 5-5. Scale bars, 1 cm. (**C**) Bin-mapping of the genomic region containing the *obv* mutation using ILs sequences. (**D**) Association plot of the GWAS for the *obv* phenotype: the x-axis represents the chromosome region where Solyc05g054030 is located, and the y-axis represents the negative log_10_ of *p*-values per SNP derived from the association analysis. The red horizontal line represents the Bonferroni-corrected genome-wide significance threshold (*p*=1.554^−08^). The boxes represent the open reading frame of Solyc05g054030. The SNP (an A-to-G transition) located in the third exon of this gene is strongly associated with the *obv* phenotype. (**E**) Gene structure of Solyc05g054030 showing exons (boxes) and introns (lines) and the position of the SNP (red arrowhead) in the third exon and DNA and protein sequence of the region around the SNP, showing that it represents a non-synonymous mutation leading to the substitution of a histidine (H) in the wild-type (WT) for an arginine (R) residue in the *obv* mutant. (**F**) Geographic distribution of accessions of wild tomatoes on the Pacific coast of Peru, Ecuador and the Galapagos Islands (inset) harboring either the WT or mutant *obv* allele. Scale bar, 100 km. (**G**) Nucleotide diversity (π) of the ancestral wild species *S. pimpinellifolium* (PIN), *S. lycopersicum* var. *cerasiforme* (CER) and big-fruit cultivars (BIG) of domesticated tomato. The horizontal lines indicates genome-wide top 5% cutoff ratio for domestication sweeps. (**H**) Frequency analysis showing the incidence of the *obv* mutation in four different categories of tomato accessions.

The presence of BSEs is not restricted to any particular phylogenetic category of plants, but rather appears to be related to functional types in the ecological succession. The distribution of BSEs in natural ecosystems is highly skewed: in a climax forest, most species from the upper strata are heterobaric, whereas those dwelling in the understory are homobaric (Kenzo et al., 2007; Inoue et al., 2015). Likewise, some crops (*e*.*g*. soybean – *Glycine max*, sunflower – *Helianthus annuus*) are heterobaric, whereas others (*e*.*g*. coffee – *Coffea* spp, cocoa – *Theobroma cacao*, typical understory species in their wild ranges) are homobaric (McClendon, 1992). What determines the distribution of these leaf functional types in natural and agricultural environments? It has been suggested that BSEs are strongly associated with higher vein densities (Baresch et al., 2019), lower resistance to water flow inside the leaf (Buckley et al., 2015) and higher vessel diameter and transpiration rates (Inoue et al., 2015). However, the lack of a suitable model system to compare heterobaric and homobaric leaf function precludes testing of these hypotheses using a genetic approach. Thus, identifying the genetic basis for BSE development would represent an important steppingstone to unveil BSE function in natural and agricultural setttings.

We previously reported that the tomato monogenic recessive mutant *obscuravenosa* (*obv*) lacks BSEs in leaves, leading to reduced water transport capacity: both stomatal conductance (*g*_s_) and hydraulic conductance (*K*_leaf_) are lower in the mutant compared to wild-type (WT) plants (Zsögön et al., 2015). Here, we identify the *OBV* gene by genetic analysis of introgression lines and genome-wide association analysis (GWAS). We find an increased frequency of the mutation in domesticated tomatoes, and analyze the geographic distribution of the mutation in the range of tomato wild relatives in South America. Furthermore, we show that *OBV* alters auxin responses and affects plant height, and leaf and fruit shape. Lastly, by analysis of *AUXIN RESPONSE FACTOR 4* (*ARF4*) loss-of-function lines, we found that a genetic module involving auxin response is involve in the development of BSEs. We discuss how this novel genetic module controlling leaf development and function could be important for ecological adaptation and for breeding selection.

## RESULTS

### Natural genetic variation for the *obv* mutation in wild tomato species

Analysis of *S. pennellii* introgression lines (ILs) in tomato showed that *OBV* locus resides on a chromosome 5 interval defined by the bin d-5E (Figure 1B and C), which contains 21 genes (Supplemental Table S1) (Jones et al., 2007; Chitwood et al., 2013). A genome wide association study (GWAS) revealed a significant single nucleotide polymorphism (SNP) in *Solyc05g054030* (Figure 1D and Supplemental Figure S1). The SNP is an A404G nucleotide change in the third exon of the gene coding sequence (CDS), resulting in a predicted histidine to arginine substitution on position 135 (H135R) of the protein (Figure 1E), which is present in different tomato cultivars harboring the *obv* mutation (Supplemental Table S2). Combining genomic and passport information we analysed the geographic distribution of the mutation in the natural range of tomato wild relatives, including the ancestral species *S. pimpinellifolium* (PIM) and the proto-domesticate *S. lycopersicum* var. *cerasiforme* (CER) found a sympatric cluster of mutant accessions in the lowlands of Ecuador and northern Peru (Figure 1F). Nucleotide diversity analysis on 360 wild and domesticated tomato accessions (available on solgenomics.net) showed that the *OBV* locus resides within a tomato domestication sweep (Figure 1G) (Lin et al., 2014). We also found that the mutation increases in frequency between accessions along the wild-domesticated continuum (Figure 1H).

### A non-synonymous single nucleotide polymorphism is responsible for the *obv* mutation

To test the hypothesis that *Solyc05g054030* encodes the *OBV* gene, we overexpressed (OE) *Solyc05g054030* with (OE^A404G^) or without (OE^A404^) the A404G polymorphism in tomato cv. Micro-Tom (MT) homozygous for the *obv* mutation (*obv/obv*) (Figure 2A). We observed complementation, *i*.*e*. correct BSE development leading to translucent veins in all *obv* plants harboring the construct lacking the A404G polymorphism (Figure 2B, and Supplemental Figure S2), but when the A404G polymorphism was present there was no complementation (Figure 2C and Supplemental Figure S3). We further confirmed that the *obv* mutant could be phenocopied by knocking down *Solyc05g054030* expression with an RNA interference (RNAi) construct (Figure 2D). We further crossed two cultivars lacking BSEs (M82 and VFN8) with either the *obv* mutant in the MT background or the *obv* mutant harboring the *OBV*^OE^ transgene. Hybrid plants derived from the former cross lacked BSEs leading to dark veins, but those derived from the latter showed phenotypic reversion, displaying transduced veins and BSEs (Figure 2E and Supplemental Figure S4). These results demonstrated that *Solyc05g054030* is *OBV*, the gene responsible for BSE development in tomato leaves, and that a single amino acid change causes the *obv* mutation, which is responsible for the switch from heterobaric to homobaric leaves in tomato. During the preparation of this manuscript the gene identity of *obv* was reported and coincides with our results here (Lu et al., 2021).

**Figure 2.**
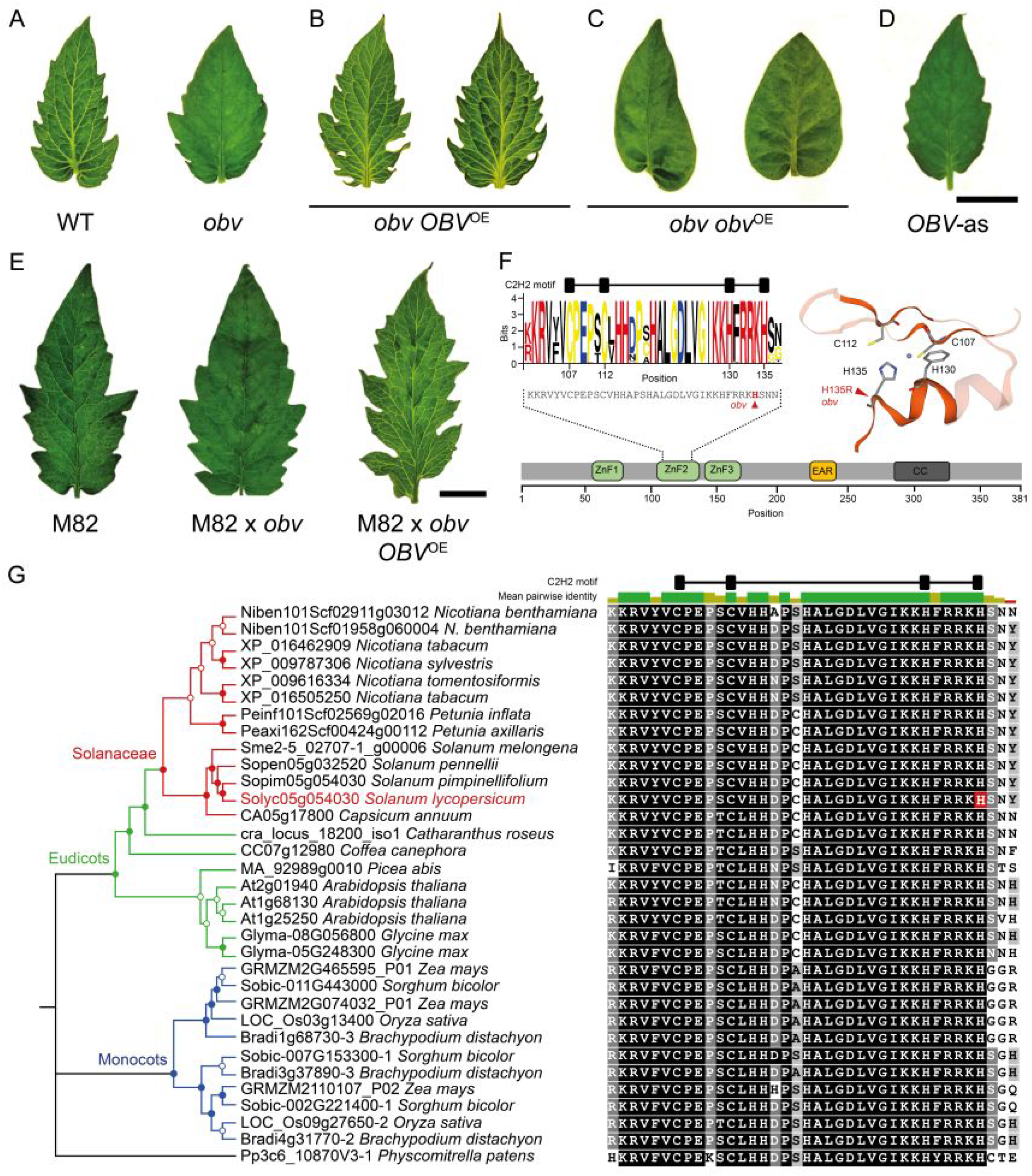
Complementation of the *obv* mutant and protein sequence analysis. (**A**) Representative terminal leaflets of tomato cultivar Micro-Tom (MT) harboring the wild-type (WT) *OBV* allele and the *obscuravenosa* (*obv*) mutant, (**B** and **C**) independent transgenic *obv* mutants overexpressing (OE) either (**B**) the WT *OBV* allele or (**C**) the mutant *obv* allele, and (**D**) an RNA interference knockdown *OBV* line in WT background. (**E**) Representative terminal leaflets of tomato cv. M82 (an *obv* mutant), an F_1_ hybrid derived from the cross between M82 and MT-*obv* and an F_1_ hybrid of M82 and transgenic MT-*obv* harboring the *OBV* overexpression transgene. Scale bars, 1 cm. (**F**) Schematic representation of the *OBV* protein showing conserved domains: ZnF (zinc finger), EAR (ethylene-responsive element binding factor-associated amphiphilic repression) and CC (coiled coil). The sequence of the second ZnF motif (ZnF2) is described as logo plot of residue conservation, with consensus sequence in the bottom. The relative sizes of letters indicate their frequency in an orthologous group of 324 proteins from 39 plant species. The total height of the letters depicts the information content of the position in bits. Right, 3D model of ZnF2 showing the relative positional arrangement of the C2H2 residues and the Zn ion ligand. (**G**) Left, maximum likelihood protein sequence tree of the *OBV* gene in selected model and crop species. Symbols on nodes represent bootstrap support values: full circles >0.75, open circles <0.75. Right, partial protein sequence alignment focussed on the second ZnF and showing the histidine (H) residue affected by the *obv* mutation in tomato (red). Shading indicates similarity to consensus (according to Blosum62 score matrix with a threshold of 1): black 100%; dark grey 80-99%; light grey 60-79%; white <60%. The bar on the top indicates the mean pairwise identity over all pairs in the column: green 100%; brown 30-99%, red <30%.

To better understand the nature of the genetic change underlying the *obv* mutation, we conducted *in silico* analyses of the gene and protein sequences of *OBV* (Figure 2F). The complete sequence of the *OBV* gene encodes a 381 amino acid protein containing three Cys_2_-Hys_2_ (C2H2) Zn finger domains, which are associated with DNA-binding capability (Persikov et al., 2015), an ethylene-responsive element binding factor-associated amphiphilic repression (EAR) domain, defined by the LxLxL motif and associated with transcriptional repression (Baile et al., 2021), and a carboxy-terminal coiled coil domain, which may function as molecular spacer or macromolecular scaffold (Truebestein and Leonard, 2016). Gene ontology (GO) terms associated with this protein are nucleic acid binding (GO:0003676) and metal ion binding (GO:0046872) (Supplemental Table S2). Using 324 protein sequences of the *OBV* orthologous group from 39 plant species (Supplemental Table S3) we pinpointed the H135R amino acid change of the *obv* mutant to a highly conserved motif of the second Zn finger domain (Figure 2G). Protein modelling showed that the C2H2 motif is contained within a ββα structure, forming a functional unit internally stabilized by chelation of a single zinc ion (Figure 2F). This suggests that the *obv* mutation leads to complete loss of protein function via disruption of its tertiary structure. Targeted phylogenetic analysis in relevant crop and model species revealed an ancient evolutionary origin of the *OBV* gene with great expansion due to duplication events (Figure 2G and Supplemental Figure S6). Protein sequence alignment of the Zn finger motif that contains the *obv* mutation in tomato showed that even the most phylogenetically distant species have high sequence identity across the domain, suggesting a conserved protein function (Figure 2G). The closest *Arabidopsis* orthologs (*INDETERMINATE DOMAIN 14, 15* and *16*), encode proteins involved in shoot gravitropism and regulation of auxin biosynthesis and transport (Supplemental Table S4 and Supplemental Figure S6) (Cui et al., 2013).

### Bundle sheath extension development is dependent on auxin signalling

We next characterized the expression profile of *OBV*, and found the gene to be highly expressed in meristems, leaf primordia, expanding and mature leaves and flowers (Figure 3A). Altered *OBV* expression levels in the transgenic lines led to changes on leaf insertion angle, leaf margin serration and fruit shape (Figure 3C-E), which hinted at a potential role for auxin as a mediator of BSE development. Auxins, like other plant hormones, exert their effect through alterations in metabolism, transport, and sensitivity (Gallei et al., 2020). We found no differences either in auxin content nor in polar auxin transport between wild type, *obv* mutant and *OBV*^OE^ lines in leaf primordia (Supplemental Figure S7). However, GUS expression driven by the *DR5* auxin-induced promoter was increased in the *obv* mutant (Figure 4A) but decreased in the *OBV*^OE^ lines (Supplemental Figure S8). Exogenous auxin also led to higher inhibition of hypocotyl elongation and stimulation of *in vitro* rhizogenesis from explants in *obv*, compared to WT or *OBV*^OE^ lines (Supplemental Figure S9). The effects were more obvious in high auxin concentrations than lower ones. This could be at least partially explained by limitations of exogenous hormone treatments, where lower concentrations are more affected by the ability of the tissue to absorb and metabolize the compounds. These results, coupled with the function of the Arabidopsis orthologs of *OBV* as mediators of auxin responses, suggested that *OBV* could be responsible for alterations in auxin signalling. Thus, we next assessed the expression profiles of *AUXIN RESPONSE FACTOR* (*ARFs*) and *AUXIN/INDOLE-3-ACETIC ACID* (*Aux/IAAs*) transcriptional regulators, which are components of auxin signal transduction (Truskina et al., 2021). We found consistent upregulation of some *ARFs* (*ARF3, 9B, 10B, 19* and *AUX*/*IAA4* and *14*) and downregulation of some *Aux/IAAs* (*Aux/IAA1A, 1B, 2, 3, 11, 13, 26* and *35*) in *OBV*^OE^ lines compared to WT (Figure 4B). In the *obv* mutant, the strongest differences were found for *ARF4* and *Aux/IAA15*, both of which were strongly upregulated compared to WT (Figure 4B).

**Figure 3.**
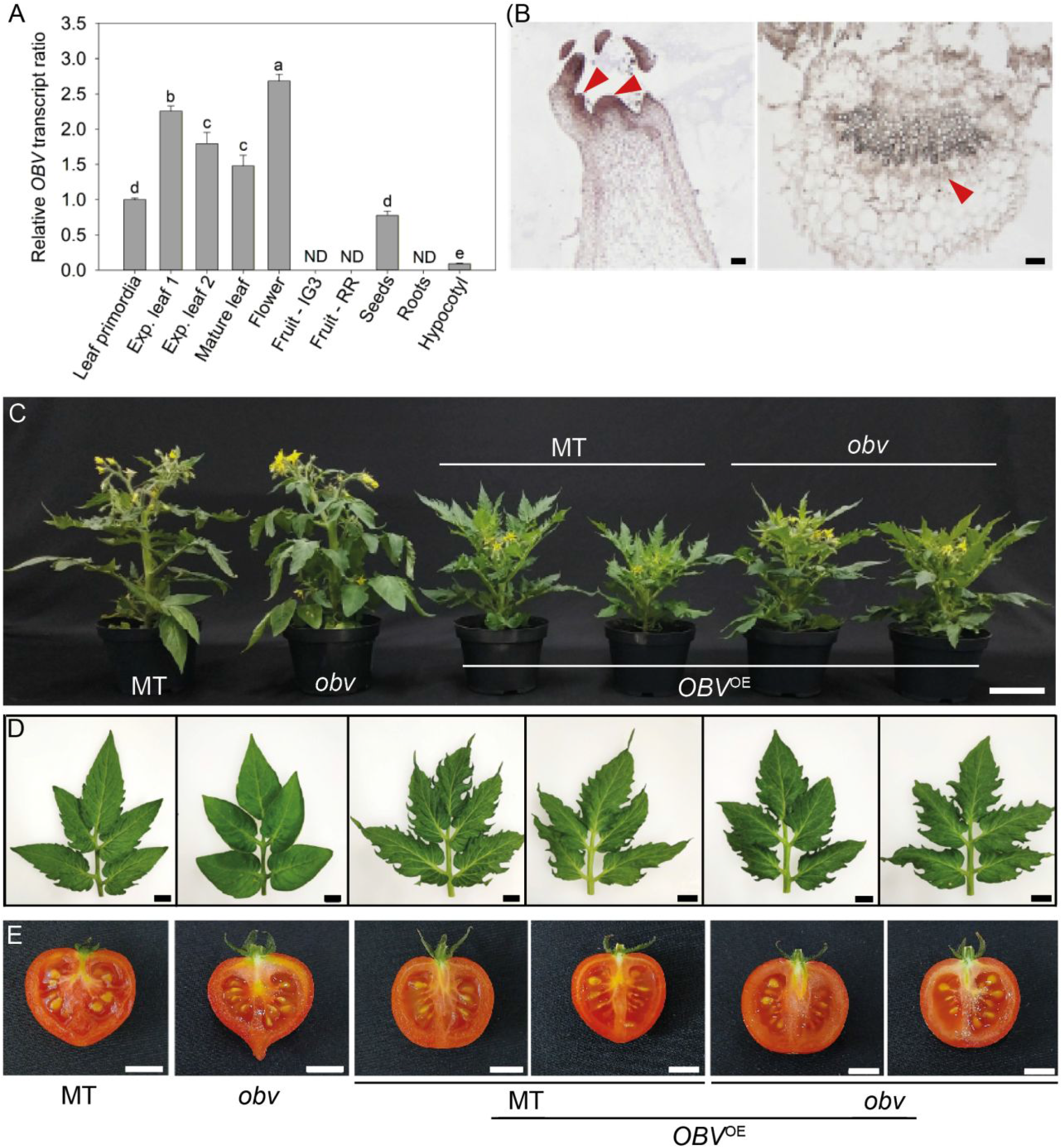
Expression pattern and pleiotropic effects of *OBV* in leaf insertion angle, leaf margin serration and fruit shape. (**A**) Relative *OBV* mRNA levels in leaves (exp. = expanding), flowers, fruits (ripening stages represented by IG3 = immature green 3 stage fruits; RR = red ripe fruits), seeds, roots and hypocotyls of tomato cultivar Micro-Tom (MT). ND = not detected. (**B**) *In situ* hybridization showing *OBV* expression patterns in a longitudinal section of the shoot apex, including the apical meristem and leaf primordia and a cross-section of an expanded terminal leaflet. Arrowheads show the specific regions where *OBV* transcripts accumulate. Scale bars = 100 µm. Representative (**C**) plants, (**D**) fully expanded fifth leaf, and (**E**) mature fruits of MT, the *obv* mutant and homozygous T_3_ transgenic lines overexpressing (OE) the *OBV* gene in either MT or *obv* mutant background. Scale bars = 10 cm (**C**) and 1 cm (**D, E**).

**Figure 4.**
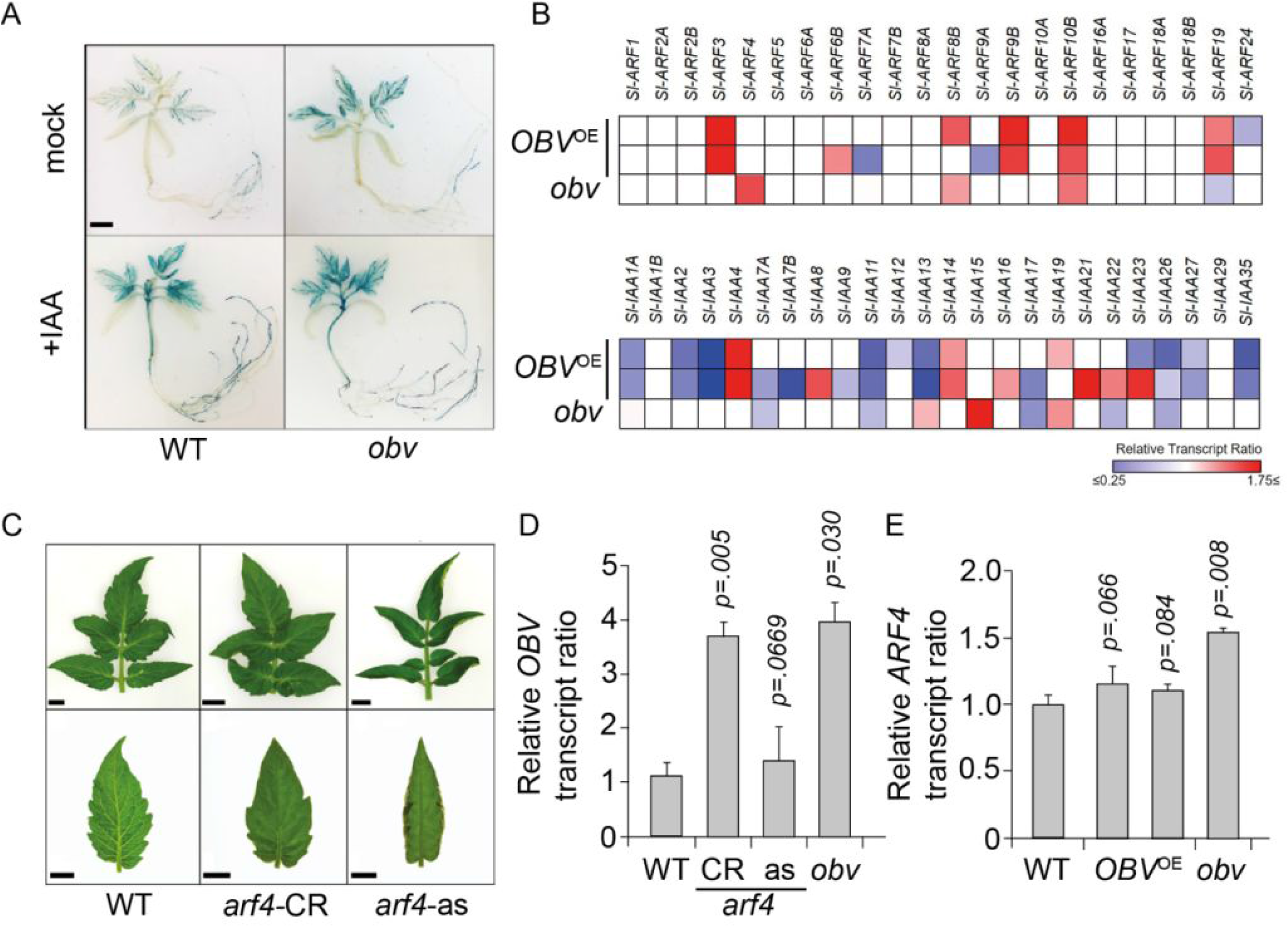
Interaction between *OBV* and auxin signalling in the control of leaf functional type. (**A**) Histochemical GUS analysis in transgenic lines harboring a homozygous *DR5*::*GUS* construct in either wild type (WT) Micro-Tom or *obv* mutant background. Seedlings were either pre-treated (+IAA) or not (mock) with 20 µM of indol-3-acetic acid. Scale bar = 1 cm. (**B**) Transcriptional profile of *AUXIN RESPONSE FACTOR* (*ARF*) and *AUXIN/INDOLE-3-ACETIC* (*Aux/IAA*) genes from leaf primordia. Heat map represents the transcript profiles in two *OBV* overexpression (OE) homozygous transgenic lines (#10 and #12) and the *obv* mutant. Values represent means of four biological replicates normalized against the corresponding WT sample. Statistically significant increases (red) or decreases (blue) in comparison with the MT are represented by colored squares (*p* <0.05). (**C**) Representative leaves and terminal leaflets of MT, a CRISPR *ARF4* mutant (*arf4*-CR) and an antisense *ARF4* knockdown line (*arf4*-as). Scale bars = 1 cm. (**D**) Relative *OBV* mRNA levels in young leaves of MT, *arf4-*CR, *arf4*-as and *obv. p*-values show significant differences to the *obv* mutant. (**E**) Relative *ARF4* mRNA levels in young leaves of MT, homozygous transgenic lines overexpressing (OE) *OBV* and the *obv* mutant. *p*-values show significant differences to the *obv* mutant.

Previous work showed that loss of *ARF4* function has strong effects on leaf development (Sagar et al., 2013; Bouzroud et al., 2020). Thus, we interrogated the potential role of *ARF4* in BSE development using a CRISPR/Cas9-generated knockout mutant (*arf4-* CR) and a transcriptionally silenced line harboring an *ARF4*-antisense (*ARF4*-as) transgene (Sagar et al., 2013). We found a lack of BSEs and the associated dark vein phenotype in leaves of both *arf4-CR* and *ARF4*-as plants (Figure 4C), which also showed the characteristic inward leaflet curling previously described (Bouzroud et al., 2020). The leaves of both *arf4-CR* and *ARF4*-as plants also showed water infiltration in the lamina and leaf margin serration patterns similar to those of the *obv* mutant (Supplemental Figure S10). Further, *OBV* expression was decreased in *ARF4-as* leaves (Figure 4D) and *ARF4* expression was increased in the *obv* mutant but restored to WT levels in the *OBV*^OE^ lines (Figure 4E). *In silico* analyses showed that the promoter region of *OBV* contains auxin-response elements (TGTCTC) (Supplemental Figure S11, Supplemental Table S4), which are typically bound by ARF proteins (Israeli et al., 2020). We conducted a dihybrid analysis to assess potential interaction between *OBV* and *ARF4* by crossing loss-of-function homozygous mutants, selfing the F_1_ hybrids and then screening visually a segregating F_2_ population. The deviation in phenotypic frequencies (Supplemental Figure 12 Supplemental Table S5), suggests that gene action between both genes is probably not independent. Taken together, these results suggest that *OBV* controls BSE development via interaction with the auxin signalling machinery.

### *OBV* has impact on CO_2_ assimilation rate and leaf hydraulic conductance

Lastly, we analysed the functional consequences of allelic variation in *OBV*. We first determined that leaf vein density (vein length per leaf area, VLA) was reduced in *obv* plants (Figure 5A, B). Vein architecture plays a key role in carbon assimilation rate and water distribution within the leaf (Sack and Scoffoni, 2013). *K*_leaf_ is a measure of how efficiently water is transported through the leaf: We found that the *obv* mutant had reduced *K*_leaf_, which was restored to wild-type levels in *OBV*^OE^ transgenic lines (Figure 5C, D). A regression analysis between VLA and *K*_leaf_ showed a strong positive correlation (*r*=0.811, *p*<0.001) between the variables, with a coefficient of determination (*R*^*2*^) of 0.657 (Figure 5E). This suggests that variation in VLA caused by *OBV* is a major determinant of variation in *K*_leaf_. Interestingly, an auxin biosynthesis mutant with reduced VLA showed reduced *K*_leaf_ and photosynthetic rate in pea (*Pisum sativum*) (McAdam et al., 2017). Net assimilation of CO_2_ (*A*_n_)-chloroplastic CO_2_ concentration curves (*C*_c_) (Figure 5E) revealed no difference in maximum Rubisco carboxylation rate (*V*_cmax_) (Supplemental Table S6) between genotypes, however, the maximum rate of light-saturated net CO_2_ assimilation (*A*_max_) was lower in the *obv* mutant (Figure 5F and Supplemental Table S6). This suggests that the presence of BSEs overrides diffusive limitations to photosynthesis through their effect on the water transport capacity of the plants (Buckley et al., 2011; Kawai et al., 2017).

**Figure 5.**
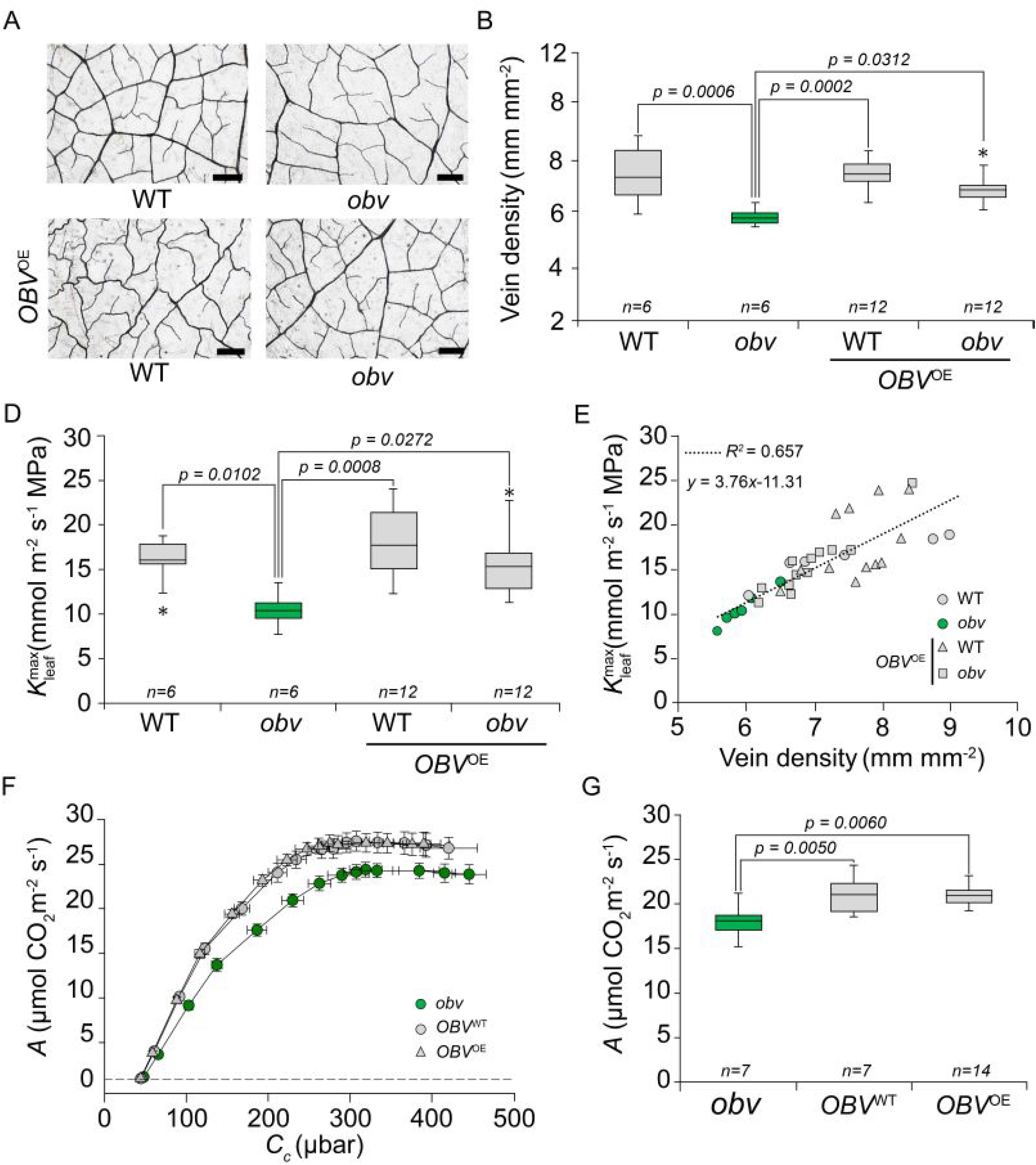
*OBV* controls vein development and leaf hydraulic conductance for hydrated leaves 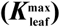. (**A**) Representative micrographs of cleared terminal leaflets of wild-type (WT) tomato cv. Micro-Tom (MT), the *obscuravenosa* (*obv*) mutant, and two independent *OBV* overexpression (OE) lines sections in either WT or *obv* background. Scale bars, 100 μm. (**B**) Leaf vein density in WT, *obv* and the transgenic lines. (**C**) values for tomato cv. M82 (*obv* mutant), a cv. M82 line harboring the wild-type *OBV* allele introgressed by conventional breeding and a transgenic *OBV*-OE line; and (**D**) in WT, *obv* and the transgenic lines. (**E**) Regression analysis of 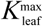and vein density for WT, *obv* and the transgenic lines. Each point corresponds to an individual measurement on a different plant. (**F**) Net photosynthetic assimilation (*A*_n_) response curve to CO_2_ concentration in the chloroplasts (*C*_c_). (**G**) Maximum assimilation rate at ambient CO_2_ and saturating irradiance derived from the curves in **F**. Boxes in box plots represent interquartile range (IQR), center line the mean, and the ends of the whisker are set at 1.5*IQR above and below the third and first quartiles, respectively. Asterisks show outliers. *p*-values for significant differences (*p*<0.05) are shown.

## DISCUSSION

Leaf BSEs fulfils significant roles in leaf function with large ecophysiological impact. Firstly, by connecting the vascular bundles to the leaf epidermis, BSEs minimize the extra-xylematic path length and favor a greater hydraulic integration in the leaf lamina (Buckley et al., 2011; Buckley et al., 2015). The stomata of heterobaric leaves may therefore operate closer to the point of embolism, while responding faster to sudden changes in xylem water potential (Zwieniecki et al., 2007; Inoue et al., 2015). This could at least partially explain the distribution of heterobaric wild tomato accessions in more arid habitats with erratic rainfall patterns (Aybar et al., 2020). Previous work has shown that environmental factors (mainly precipitation regime and intensity of light competition) predict adaptive morphological differentiation between wild tomatoes in their native range (Nakazato et al., 2008; Nakazato et al., 2010). The most dramatic changes occur in the transition from PIM to CER, which have marked range differences, and include increased leaf area and leaf water content but faster wilting under drought in the latter (Nakazato et al., 2008). Secondly, photosynthetic assimilation rates are higher in heterobaric leaves due to the optimization of light transmission within the leaf lamina (Nikolopoulos et al., 2002). BSEs can function as ‘transparent windows’ that enrich neighbouring mesophyll cells (Karabourniotis et al., 2000), or, in the case of C_4_ plants, the bundle sheath itself, with high levels of photosynthetically active radiation (400-700 nm) (Bellasio and Lundgren, 2016). The higher photosynthetic assimilation rate in plants harbouring the functional *OBV* allele support the contention that heterobaric and homobaric leaves differ in their ‘carbon-gain strategy’ (Liakoura et al., 2009). Lastly, the presence of BSEs increases the plasticity of minor vein density in response to growth light intensity, adjusting water supply and photosynthetic rate to specific environmental conditions (Barbosa et al., 2019). The sum of these effects on leaf function suggests that the presence of BSEs alters the mechanisms that produce the key relations of the leaf economic spectrum (Wright et al., 2004), and is adaptive in local habitats by realizing a trade-off between construction costs and functional gains. This raises the question of what the main driver for the loss of BSEs in domesticated tomato cultivars could be (Jones et al., 2007). One possible explanation it that the heterobaric/homobaric ecological distribution logic was broken down by the passage to an agricultural setting and, in this case, the homobaric phenotype is functionally more advantageous due a trade-off with the structural and functional costs of BSEs (Baresch et al., 2019). Furthermore, since the *obv* mutation increases sensitivity to auxin, we cannot exclude the selection of a favorable pleiotropic effect of this hormone on plant development and productivity (Hu et al., 2018).

The role of auxin in the regulation of leaf morphogenesis and development of vascular tissue is well described (Vanneste and Friml, 2009). Auxin controls the leaf margin dissection, which can vary between round/ovate and serrated (‘toothed leaves’) (Koenig et al., 2009). Loss of *OBV* function leads to rounder leaves, whereas overexpression increases leaf dissection and margin serration. In woody species, increased margin dissection has been associated with colder climates (McKee et al., 2019), however it is not known if this association is extensive to herbaceous species like tomato and its wild relatives. Thus, further analysis of our genotypes could provide novel insights on the functional significance of leaf margin serration in annual herbs. Auxin is also a key controller of fleshy fruit development, and plays a role in fruit set upon fertilization and in determination of final fruit size through control of cell division and expansion (Fenn and Giovannoni, 2021). Loss of *ARF4* function leads to a ‘heart-shaped’ fruit (see Figure 2D in Sagar et al., 2013), which is phenocopied by the *obv* mutant. Here we also showed that *arf4* mutants lack BSEs, which coupled to the reciprocal inhibition of gene expression between *ARF4* and *OBV*, and their control of BSE development, suggest that they may be operating together at some level. Future work will address potential physical protein-DNA and protein-protein interactions. Our results showed that the H135R substitution impairs *OBV* function, and suggest that it may affect tridimensional protein structure.

The strength of selection against a given amino acid replacement is a function of the chemical similarity between the original amino acid and the non-synonymous one (Yampolsky et al., 2005). In *obv*, the mutation in position 135 replaces a histidine for an arginine. While both are polar amino acids, histidine is unique with regard to other chemical properties, which means that is does not substitute particularly well with any other amino acid. Our modellling results showed that H135 most likely participates in a metal binding site, acting together with cysteines or other amino acids. It is thus notable that the H135R is not only highly conserved but also found in higher frequency in modern tomato cultivars than in its wild relatives. Homobaric (*i*.*e*. homozygous *obv* mutants) accessions of the tomato wild relatives are confined to the lowlands of the coastal regions of Ecuador and northern Peru, which show a milder, more humid climate (Ilbay-Yupa et al., 2021). These conditions change rapidly going south, due to decreased rainfall, or west, due to increasing altitude in the Andes Mountain range (Moyle, 2008; Aybar et al., 2020). Thus, either purifying selection against the mutation is relaxed under milder conditions, or positive selection favours the homobaric leaf type, selecting against heterobaric leaves’ higher construction or operational cost (Read and Stokes, 2006). In wild PIM populations a combination of purifying and balancing selection maintains polymorphism in plant resistance genes (Caicedo, 2008).

Three key points to consider when attempting to infer the dynamics of the preservation of a polymorphism in a species are recombination rates (Roselius et al., 2005), mating system (Glémin, 2007) and effective population size (Gaut et al., 2018). Recent work has shown a generally conserved recombination landscape between tomato and its wild relatives (Fuentes et al., 2021). As for mating system, tomato domestication was accompanied by a transition from allogamous wild relatives to autogamous domesticates (Benoit, 2021). Notably, self-fertilization increases the efficiency of purifying selection against slightly deleterious recessive mutations (Byers and Waller, 1999) but aids fixation of favorable recessives (Charlesworth, 1992). Lastly, in species with small effective population size, or that have undergone episodes of strong population bottlenecks, like tomato during its domestication (Razifard et al., 2020; Razifard et al., 2021), slightly deleterious mutations can increase in frequency, and eventually even be fixed due to drift (Whitlock, 2000). The strong developmental and functional effects of *OBV*, the skewed geographical distribution of the mutation, and its increasing frequency along the continuum of wild-to-domestic species suggest, however, that genetic drift alone may not be the most satisfactory explanation. An alternative possibility is that the changes in *OBV* allele frequency may have been caused by ‘genetic hitchhiking’, *i*.*e*. selection on a closely linked locus.

The *OBV* locus resides relatively close (∼70 kpb) to *SELF PRUNING 5G* (*SP5G*), a repressor of flowering in long days (Soyk et al., 2017). A hypomorphic allele of this gene was under strong artificial selection during domestication: A severely mitigated long-day induction of *SP5G* caused by a 52-bp deletion explains the nearly complete loss of daylength-sensitive flowering in domesticated tomato (Zhang et al., 2018; Song et al., 2020). Previous work had suggested that *obv* frequency had increased in processing tomatoes due to selection on *SP5G* for improved growth habit (Jones et al., 2007). It was subsequently shown that *SP5G* is located within a ‘domestication sweep’ as revealed by a drastic reduction of nucleotide diversity between PIM and CER (Lin et al., 2014). However, analysing sequence data from 72 wild and domesticated tomato accessions, we did not find evidence of significant linkage disequilibrium between the *sp5g* and *obv* mutations (Supplemental Tables S7 and S8), suggesting that selection for day neutrality or growth habit (both controlled by *sp5g*) was not a driver of BSE loss through genetic hitchhiking (Barton, 2000). Further, even though sequence variation on chromosome 5 is the main driver of the divergence between fresh and processing tomatoes (Robbins et al., 2011), the incidence of *obv* is very similar in both categories of cultivars. Lastly, a drastic change in the proportion of non-synonymous mutations occurred during the transition from PIM to CER, and many neutral or beneficial non-synonymous variants were selected during the northward spread of the latter (Razifard et al., 2021). Thus, the population dynamics of the *obv* mutation ought to be considered within this new evolutionary framework. While it remains an open question whether homobaric leaves are advantageous in an agricultural setting, our results suggest that this may be the case, and the genotypes described here are well-suited to conduct functional analyses of each leaf type under contrasting environments. Analysis of natural genetic variation could also address the ecological contribution of BSEs to adaptation.

## CONCLUSION

Our findings represent an entry point to unravel leaf functional design through a gene-focused approach and provide a molecular anchor for the analysis of phenotypic co-variation in leaf anatomical and physiological traits. They also suggest that the divergence in leaf functional types results from a trade-off conditioned by environmental factors and may have adaptive value. The conservation of *OBV* in angiosperms could be leveraged to explore its roles in other plant major crop species and potentially tailor different leaf types to specific agronomic settings. For instance, BSEs were recently shown to provide a pathway for P uptake following foliar fertilization in spring barley (*Hordeum vulgare*) (Arsic et al., 2020). Thus, our discovery of a novel genetic module controlling the switch between homobaric and heterobaric leaves could have important ecological implications and agricultural applications.

## MATERIALS AND METHODS

### Plant material, growth conditions and breeding

Tomato seeds (*Solanum lycopersicum* cv. Micro-Tom, MT and cv. M82) were sown on polyethylene trays containing Tropostrato^®^ commercial substrate (São Paulo, Brazil) and grown in a greenhouse in Viçosa (642 m asl, 20°45’S; 42°51’W), with an average irradiance of ∼800 µmol m^−2^ s^−1^, photoperiod 12/12-h and air temperature 26/18°C day/night. Upon appearance of the first true leaf, seedlings of MT and M82 were transplanted to pots with a capacity of 350 mL and 3000 mL, respectively. Soil was fertilized with 2 g L^−1^ NPK (10-10-10) and 4 g L^−1^ limestone. For *in vitro* cultivation, seeds were sown in flasks containing 30 mL of half-strength MS medium gellified with agar 4 g L^−1^, pH 5.7 ± 0.05. The seedlings were kept under controlled conditions: photoperiod of 16-h/8-h day/night, light intensity of 45 ± 3 μmol m^−2^ s^−1^ and temperature of 25 ± 1 °C.

### Identification of *OBV* homologs and phylogenetic inference

To identify *OBV* homologs in tomato and other plant species, we retrieved the ortholog group of proteins using the Plant Transcription Factor Database (planttfdb.gao-lab.org) (Supplemental Table S9). We next aligned the sequences from the model species *A. thaliana* with MUSCLE. The alignment was submitted to trimAl for alignment trimming and then submitted to FastTree for tree inference. Trees were visually inspected using FigTree (tree.bio.ed.ac.uk/software/figtree/). Further phylogenetic inference using only *OBV* homologs were performed using MUSCLE, for sequence alignment, trimAl, for alignment trimming, SMS, for evolutionary model selection, and PHYML, for maximum likelihood tree inference. Final trees were annotated with taxonomic information from NCBI Taxonomy using TaxOnTree (bioinfo.icb.ufmg.br/taxontree). The plant proteomes used in this work were retrieved from Sol Genomics Network (solgenomics.net), for tomato (ITAG v.4.0), and from Uniprot (www.uniprot.org), for other species.

### Gas exchange analyses

Gas exchange analyses were performed in adult plants of cv. M82, M82-*OBV* (cv. M82 harboring the functional *OBV* allele) and M82-*OBV*^OE^ (M82 plants harboring an *OBV* overexpression construct). All the evaluations described below were measured in terminal leaflets of the fifth expanded leaf. Gas exchange parameters were determined simultaneously using an open-flow infrared gas analyzer (IRGA) system (model LI-6400XT, Li-Cor Inc., Lincoln, NE, EUA). The equipment was configured to provide a light intensity of 1000 μmol m^−2^ s^−1^, CO_2_ concentration of 400 μmol mol^−1^, with the air flow in the chamber regulated to 300 μmol s^−1^. The *A*/*C*_i_ response curves were measured under ambient O_2_ and temperature, using PAR of 1000 μmol m^−2^ s^−1^ and injection of incremental CO_2_ concentrations into the chamber (50, 100, 200, 300, 400, 500, 600, 700, 800, 900, 1000, 1200, 1400, 1600, 1800 µmol mol^−1^). Calculations of chloroplast concentrations of CO_2_ (*C*_c_) and mesophyll conductance (*g*_m_) were performed using the Harley method. The CO_2_ compensation point quantified previously for tomato was used as reference to calculate *g*_m_ and *C*_c_.

### Statistical analysis

The experimental design was completely randomized. Data were submitted to analysis of variance (ANOVA) and the means were compared by Tukey’s test at 5% level of significance (*P* ≤ 0.05).

## Supplemental data

### Materials and Methods

Supplemental Figure 1. Mapping and identification of the *OBV* candidate gene

Supplemental Figure 2. Complementation of the *obv* mutant with the OBV functional allele Supplemental Figure 3. Complementation of the *obv* mutant and knockdown of the *OBV* gene

Supplemental Figure 4. Morphology of OBV overexpressing plants in hybrid M82 × MT and VFN8 × MT backgrounds

Supplemental Figure 5. Maximum likelihood tree of the OBV family subclade compromising tomato OBV

Supplemental Figure 6. Phylogenetic reconstruction of the OBV protein family in tomato, *Arabidopsis* and *Physcomitrella patens*

Supplemental Figure 7. Influence of OBV on leaf free auxin concentration and polar auxin transport in hypocotyl explants

Supplemental Figure 8. Reduction of auxin sensitivity in OBV-overexpressing lines revealed by histochemical GUS assays

Supplemental Figure 9. Hypocotyl elongation and in vitro rhyzogenesis assays show that OBV alter auxin sensitivity

Supplemental Figure 10. Control of BSEs development by ARF4 and OBV

Supplemental Figure S11. Phenotype of representative leaves from a dihybrid *arf4* × *obv* cross

Supplemental Figure S12. *In silico* analysis of the *OBV* (*Solyc05g054030*) promoter region. Supplemental Table S1. *OBV* candidate genes contained in bin d5-E of the *S. pennellii* introgression lines

Supplemental Table S2. *OBV* alleles in different *S. lycopersicum* accession and wild species Supplemental Table S3. Enrichment analysis by Gene Ontology for the *OBV* gene Supplemental Table S4. Analysis of the *OBV* promoter

Supplemental Table S5. Contingency table and goodness of fit calculation for *obv* and *arf4* phenotpyic segregation ratios

Supplemental Table S6. Gas exchange and chlorophyll fluorescence parameters determined in fully-expanded leaves of tomato cv. M82 (*obv* mutant), complemented M82-*OBV* and *OBV*-overexpressing plants

Supplemental Table S7. Statistical analysis of linkage disequilibrium between *OBV* and *SP5G* in 72 tomato accessions.

Supplemental Table S8. Sequences used for phylogenetic and alignment analyses

Supplemental Table S9. Oligonucleotide DNA sequences for PCR primers used in this study

Supplemental Table S10. Accessions used for the linkage disequilibrium analysis between *OBV* and *SP5G*

## Acknowledgements

We thank Profs Alisdair Fernie and Ralph Bock (Max-Planck Institute, Germany), Prof José Jiménez-Gómez (INRA Versailles, France), Prof. Andrew J. Thompson (Cranfield University) and Diego S. Reartes for valuable discussions and input on the manuscript. We also gratefully acknowledge the support and contributions of the UFV Plant Physiology Graduate Program.

## Notes

### Competing Interest Statement

The authors have declared no competing interest.

### Summary of Updates

Manuscript revised and expanded

