## Supplemental Figures for "Auxin-driven ecophysiological diversification of leaves in domesticated tomato"

Supplementary Materials and Methods

Figures S1 to S11

References

### Materials and Methods

#### Plant material, growth conditions and breeding

Tomato seeds (*Solanum lycopersicum* cv. Micro-Tom, MT and cv. M82) were sown on polyethylene trays containing Tropostrato® commercial substrate (São Paulo, Brazil) and grown in a greenhouse in Viçosa (642 m asl, 20º45´S; 42º51´W), with an average irradiance of ~800 µmol $m^{-2}$ $s^{-1}$, photoperiod 12/12-h and air temperature 26/18°C day/night. Upon appearance of the first true leaf, seedlings of MT and M82 were transplanted to pots with a capacity of 350 mL and 3000 mL, respectively. Soil was fertilized with 2 g $L^{-1}$ NPK (10-10-10) and 4 g $L^{-1}$ limestone. For *in vitro* cultivation, seeds were sown in flasks containing 30 mL of half-strength MS medium (Murashige and Skoog, 1962) gellified with agar 4 g $L^{-1}$, pH 5.7 ± 0.05. The seedlings were kept under controlled conditions: photoperiod of 16-h/8-h day/night, light intensity of 45 ± 3 µmol $m^{-2}$ $s^{-1}$ and temperature of 25 ± 1 ˚C.

The tomato cv. M82 that harbours the recessive *obv* mutant allele was used to generate M82 near-isogenic lines carrying the *OBV* functional allele (M82-*OBV*). We performed the introgression of both the wild-type *OBV* allele and the $OBV^{OE}$ transgene from wild-type MT plants or MT harbouring an $OBV^{OE}$ transgene, following the scheme illustrated in Supplemental Figure S12. Briefly, plants were grown in greenhouse under conditions described above. M82 and MT were crossed using the former as female parental and the latter as pollen donor. $F_1$ plants were self-pollinated generating a segregating $F_2$ progeny. The $F_2$ seedlings resulting were screened for translucent veins and the bigger stature of M82. Selected plants were backcrossed (BC) with M82 parental, producing $BC_1F_2$ seeds. This procedure of crossing and screening was carried until the $BC_3$ generation, where plants were self-pollinated and the progeny screened for homozygous plants ($BC_3F_n$).

#### GWAS analysis

The online repository of Solanaceae genetics and genomics information Sol Genomics Network (solgenomics.net) was used to access the database of 360 genomes of tomato cultivars and wild species (Lin et al., 2014). We focussed on a discrete region of chromosome 5 based on the previously mapped location of *OBV* (Jone et al., 2007) and analysed genetic variations present in all 37 genes contained in the bin d-5E (Supplemental Table S1). We then used the extensive database from the Tomato Genetics

Resource Center, TGRC (tgrc.ucdavis.edu) to retrieve phenotypic data for different accessions of tomato cultivars and wild species and conduct the genotype × phenotype association.

The Genome-Wide Association Study (GWAS) was performed using data of 37 tomato accessions (25 wild-type and 12 mutants for *obv* phenotype, Supplemental Table S2) and the Plink (v.1.07) software (Purcell et al., 2007). A total of 3,218,302 single nucleotide polymorphisms (SNPs) with a minor allele frequency (MAF) greater than 5% were used for association analysis. The whole-genome significant cutoff was defined as P=1.554e-08 based on the Bonferroni correction method. A Manhattan plot was generated using the R package qqman (Turner, 2014). Gene annotation was obtained from International Tomato Annotation Group (ITAG, v.2.4) and visualized using the R package Gviz (Hahne and Ivanek, 2016).

Structural modeling analysis

The analyses of functional domains in the OBV protein were done using the functional domain prediction tool from Interpro (www.ebi.ac.uk/interpro/), and the Logo plot was obtained by comparing of 324 protein sequences from the ortholog group of 39 plant species, using EVcouplings (evcouplings.org). The 3D model of the wild-type OBV protein was generated on SwissModel (swissmodel.expasy.org).

*OBV* homologs identification and phylogenetic inference

To identify *OBV* homologs in tomato and other plant species, we retrieved the ortholog group of proteins using the Plant Transcription Factor Database (planttfdb.gao-lab.org). We next aligned the sequences from the model species *A. thaliana* with MUSCLE (Edgar, 2004). The alignment was submitted to trimAl (Capella-Gutiérrez et al., 2009) for alignment trimming and then submitted to FastTree (Price et al., 2010) for tree inference. Trees were visually inspected using FigTree (tree.bio.ed.ac.uk/software/figtree/). Further phylogenetic inference using only *OBV* homologs were performed using MUSCLE, for sequence alignment, trimAl, for alignment trimming, SMS (Lefort et al., 2017), for evolutionary model selection, and PHYML (Guindon et al., 2010), for maximum likelihood tree inference. Final trees were annotated with taxonomic information from NCBI Taxonomy using TaxOnTree (bioinfo.icb.ufmg.br/taxontree). The plant proteomes used in this work were retrieved from Sol Genomics Network, for tomato (ITAG v.4.0) (Hosmani et al., 2019), and from Uniprot (www.uniprot.org), for other species.

Molecular cloning

The coding sequence of *OBV* gene (1,149 pb) was reverse transcribed and amplified by PCR (Polymerase Chain Reaction) from RNA extracted from leaves of *Solanum lycopersicum* cv. Micro-Tom (wild-type) or the *obv* mutant, using the primers listed in Supplemental Table S10. Young leaves (~2.0 cm) were collected for extraction of total RNA using Trizol® (Ambion, Life Technology), according to the manufacturer's recommendations. The cDNA synthesis was performed with SuperScript™ III First-Strand Synthesis System (Invitrogen). The amplicons were purified and cloned independently in the plasmids PCR8/GW/TOPO® (Invitrogen). Subsequently, each fragment was combined in the overexpressing vector pK7WG2D,1 through Gateway® technology (Karimi et al., 2002). Expresion of the gene of interest is driven by a 35S promoter. The correct assembly of the construct was confirmed by PCR and Sanger sequencing.

For post-transcriptional gene silencing we used an interference RNA (RNAi) construct. A specific fragment of 223 pb (Appendix 1) from coding sequence of *OBV* gene was selected using BLAST queries against the Sol Genomics database (solgenomics.net/, ITAG release 2.40). The amplified fragment was purified and cloned into the plasmids PCR8/GW/TOPO® (Invitrogen), and then combined in the gene silencing vector pK7GWiWG2 (I) (Karimi et al., 2002). The assembly was confirmed by PCR and Sanger sequencing. The confirmed plasmids for overexpression or silencing were used to transform *Agrobacterium tumefaciens* (EHA105) by heat shock as previously described (Pino et al., 2010).

Plant transformation

*A. tumefaciens* EHA105 harbouring the plasmids described above was used to perform tomato plants transformation as described (Pino et al., 2010). Briefly, seeds were sterilized by agitation in 30% (v/v) commercial bleach (2.7% w/v sodium hypochlorite) for 15 min and rinsed with distilled water. 30 seeds of Micro-Tom or *obv* mutant were inoculated into flasks containing half-strength MS medium (Murashige and Skoog, 1962), vitamins B5, sucrose 30 g $L^{-1}$ and agar 4 g $L^{-1}$. The pH of the medium was adjusted to 5.7

± 0.05 and sterilized by autoclaving. The seeds were kept four days in the dark, 25 ± 1 °C, followed by four days under photoperiod of 16-h/8-h day/night and intensity 45 ± 3 µmol $m^{-2}$ $s^{-1}$. The cotyledons of 8-days-old seedlings were sectioned and used in co-cultivation with *A. tumefaciens* in Root Inducing Medium (RIM) for 2 days, 25 ± 1 °C in the dark. Then, the explants were transferred to Shoot Inducing Medium (SIM) and were kept in 25 ± 1 °C under photoperiod of 16-h/8-h and light intensity of 10-20 µmol $m^{-2}$ $s^{-1}$, for four weeks. The explants were recultivated to fresh medium twice during this period. Well-developed shoots (≥ 5 cm) were transferred to flasks containing hormone-free MS medium, for shoot elongation and rooting. At all *in vitro* stages, the medium was supplemented with 300 mg $L^{-1}$ timentin to suppress *A. tumefaciens* growth and 100 mg $L^{-1}$ kanamycin for *in vitro* selection of transgenic plants.

Selection of overexpressing and silencing lines

Regenerated overexpressing and silencing $T_0$ plants were acclimatized under controlled conditions, photoperiod 16-h/8-h day/night, irradiance of 150 µmol $m^{-2}$ $s^{-1}$ and temperature of 25 ± 1 °C, using autoclaved Tropostrato® substrate and vermiculite (1:1) for four weeks. Subsequently, $T_0$ transgenic plants were transferred to greenhouse for genotyping and seed production.

To verify the transgene insertion, genomic DNA was extracted from young leaflets (Edwards et al., 1991). First, the presence of the resistance gene *NEOMYCIN PHOSPHOTRANSFERASE II* (*neo*) was confirmed by PCR, for both constructs. A second confirmation was made to identify the transgene using a forward primer for *pCaMV 35S* and reverse specific for the construct of interest. The primers used are listed in Supplemental Table S10.

To obtain homozygous plants, overexpressing and silencing lines were manually self-pollinated producing $T_1$, $T_2$, $T_3$ and $T_4$ seeds. Progeny tests were carried out on seeds derived from single plants through foliar spray of antibiotic (kanamycin 400 mg $L^{-1}$ for five consecutive days). The plant populations (n=30) with 100% plants antibiotic resistant (assessed visually as absence of leaf yellowing following kanamycin application) were considered homozygous. The assay was repeated on at least two subsequent generations.

Real-time quantitative PCR (RT-qPCR)

To assess the *Aux/IAA* transcriptional profile, leaf primordia (≤ 0.5 cm) were collected from Micro-Tom (wild-type), *obv* mutant and overexpressing lines. RNA was purified using ReliaPrep™ RNA Miniprep Systems (Promega), according to the manufacturer recommendations. The cDNA synthesis was performed with SuperScript™ III First-Strand Synthesis System (Invitrogen). Real-Time quantitative PCR reactions (qRT) were performed in a thermocycler Real-Time StepOnePlus PCR (Applied Biosystems) with a final volume of 14 µL using reagent SYBR Green Master Mix (Thermo Fisher Scientific). The melting curves were analyzed for nonspecific amplifications and dimerization of primers. Absolute fluorescence data were analyzed using the software LinRegPCR (Ruijter et al., 2009) to obtain the values of quantification cycle (Cq) and calculate the primer efficiency. The abundance of transcripts was normalized against the geometric mean of two reference genes, *TIP4* and *EXPRESSED* (Expósito-Rodriguez et al., 2008).

To analyse the *OBV* expression profile in wild-type MT plants, the following tissues were collected: germinated seeds, root tips (≤ 2 cm), hypocotyls, leaf primordia (≤ 0.5 cm), expanding leaf 1 (≤ 2 cm), expanding leaf 2 (≤ 3 cm), mature leaf, flowers (petals + anthers), immature green 3 stage fruits (IG3) and red ripe fruits (RR). RNA extraction and analyses was conducted as described above. The sequences of all primers used in this study are listed on Supplemental Table S10.

*In situ* hybridization

The *OBV* expression analyzed by *in situ* hybridization assay was performed as described previously (Oliveira et al., 2017). Apical meristems and leaf primordia of wild-type MT plants were collected and processed.

Phenotypic characterization

The morphological characterization was made in MT, *obv* mutant and overexpression lines in $T_3$ homozygous transgenic plants, after anthesis. Plant height was measured from the soil level to the first inflorescence. The length and diameter of the internodes were obtained by measuring the fourth, fifth and sixth internodes counted from the bottom up. Stem diameter was measured from the cotyledons level using a mechanical pachymeter (Mitutoyo® Vernie Caliper model, Japan). Leaf number to the first inflorescence was obtained by counting of the leaves on main stem, from the bottom up. The leaf angle was determined based on the insertion of the fifth, sixth and seventh leaf, using a protractor.

Biomass accumulation was obtained through root, stem and leaves dry weight by destructive analysis and oven-drying 60 days after germination. All measurements were conducted in at least 12 plants per genotype.

For leaf characterization, three terminal leaflets were selected per plant, in eight plants per genotype. The leaflets were digitized using an HP Scanjet G2410 scanner (Hewlett-Packard, Palo Alto, California, USA). Total leaf area was obtained digitizing all leaves and calculating the area in Image-Pro Plus. Specific leaf area (SLA) was calculated through the ration between leaf area (LA) and leaf dry weight (LDW), as described by the equation: SLA ($cm^2$ $g^{-1}$) = LA/LDW.

<u>Gas exchange, chlorophyll *a* fluorescence and *A*/*Ci* curves</u>

Gas exchange analyses were performed in adult plants of cv. M82, M82-*OBV* (cv. M82 harbouring the functional *OBV* allele) and M82-$OBV^{OE}$ (M82 plants harbouring an *OBV* overexpression transgene). All the evaluations described below were measured in terminal leaflets of the fifth expanded leaf. Gas exchange and chlorophyll *a* fluorescence parameters were determined simultaneously using an open-flow infrared gas analyzer (IRGA) system (model LI-6400XT, Li-Cor Inc., Lincoln, NE, EUA) equipped with an integrated fluorescence chamber of 2$cm^2$ (6400-40 Leaf Chamber, Li-Cor Inc.). The equipment was configured to provide a light intensity of 1000 μmol $m^{-2}$ $s^{-1}$, $CO_2$ concentration of 400 μmol $mol^{-1}$, with the air flow in the chamber regulated to 300 μmol $s^{-1}$. For evaluation of dark respiratory activity ($R_d$), a 6 $cm^2$ leaf chamber with air flow set at 100 μmol $s^{-1}$ was used. The effective quantum yield of photosystem II ($\phi PSII = (Fm'–Fs)/Fm'$) was determined by measuring the steady-state fluorescence (*Fs*) and the maximum fluorescence (*Fm'*), applying a saturating light pulse of 8000 μmol $m^{-2}$ $s^{-1}$ photons (Genty et al., 1989). Photochemical quenching ($qP = (Fm'–Fs)/(Fm'–F0')$) was calculated as described (Oxborough and Baker, 1997). The apparent rate of electron transport ($ETR = 0.5 \times 0.84 \times \phi PSII \times$ PAR) was calculated as described in (Melis et al., 1987), where 0.5 = fraction of the excitation energy distribution in the FSII; 0.84 = fraction of light absorbed by leaves; and PAR = photosynthetically active radiation.

The $A/C_i$ response curves were measured under ambient $O_2$ and temperature, using PAR of 1000 μmol $m^{-2}$ $s^{-1}$ and injection of incremental $CO_2$ concentrations into the chamber (50, 100, 200, 300, 400, 500, 600, 700, 800, 900, 1000, 1200, 1400, 1600, 1800 μmol $mol^{-1}$). Calculations of chloroplast concentrations of $CO_2$ ($C_c$) and mesophyll conductance ($g_m$) were performed using the Harley method (Harley et al., 1992). The $CO_2$

compensation point quantified previously for tomato (Muir et al., 2017) was used as reference to calculate $g_m$ and $C_c$. The maximum Rubisco carboxylation velocity ($V_{cmax}$), the maximum capacity for electron transport rate ($J_{max}$), and the use of phosphate trioses (TPU) were estimated based on $C_i$ and $C_c$ and normalized to 25°C (Long and Bernacchi 2003). Stomatal, mesophyll and biochemical limitations were calculated as described (Grassi and Magnani, 2005). $CO_2$ and water vapor leaks were corrected according to the methodology described in (Rodeghiero et al., 2007).

Leaf hydraulic conductance ($K_{leaf}^{max}$) determinations

Leaf water potential ($\Psi_{leaf}$) was measured in the terminal leaflet of the fifth expanded leaf of MT, *obv* mutant and transgenic overexpression lines *OBV*$^{OE}$, using a Scholander-type pressure chamber (model 1000, PMS Instruments, Albany, NY, USA). The leaf hydraulic conductance ($K_{leaf}^{max}$) was estimated using the transpiration rates and the water potential difference between the transpiring and non-transpiring leaflet. The non-transpiring leaflet consisted of the lateral leaflet of the same leaf, which was covered with plastic film in the night before the measurements. $K_{leaf}^{max}$ was calculated according to equation:

$$K_{leaf}^{max} = E / (\Psi_L - \Psi_X)$$

Where $E$ is the transpiration rate (mmol $m^{-2}$ $s^{-1}$) determined during gas exchange measurements, and ($\Psi_L$ – $\Psi_X$) corresponds to the pressure gradient between the transpiring and non-transpiring leaflet (MPa) (Sack and Holbrook, 2006). The parameters were determined in six plants per genotype.

Anatomical analyses

Expanded terminal leaflet from the fifth leaf were used for anatomical analyses. To obtain cross sections, a 1 × 0.5 cm fragment was excised from the leaf lamina and fixed in FAA70 (70% formalin-acetic acid-alcohol) for 24 h. Subsequently, the samples were dehydrated in an ethanol series (70%, 85% and 95%), under vacuum (-20 polHg). Pre-infiltration was performed with resin and ethanol 95% (1:1) for 2 h under vacuum (-20 polHg), and infiltration was done in resin for one week. Cross-sections of 5 µm thickness were mounted on blades and stained with 0.05% toluidine blue. The images were analyzed under a light microscope (Zeiss AxioScope A1, Jena, Germany).

Vein density measurements

Vein density was measured in six terminal leaflets per genotype. The leaflets were collected and fixed in methanol 95% for 48 h, then the solution was changed to lactic acid 100% and incubated at 100°C until complete clarification (5-6 hours). The diaphanized leaflets were analyzed using a light microscope (Zeiss AxioScope A1, Jena Germany) and vein density was measured as length of veins per unit leaf area, using the software Image Pro-Plus® (imagej.nih.gov/ij/).

Histochemical GUS analysis

Seedlings of 20-days-old MT, *obv* mutant and overexpression lines were crossed to transgenic MT or *obv* lines carrying the auxin-responsive synthetic promoter (*DR5*) (Ulmasov et al., 1995) fused to the reporter gene *UID* (encoding β-glucuronidase). $F_1$ hybrid seeds were harvested and grown *in vitro* as described above. For each genotype, four 3-week-old seedlings were incubated in exogenous auxin (20 µM indole-3-acetic acid, IAA), or mock treated for 3 h. The seedlings were incubated at 37 °C for 12 h in GUS staining solution (100 mM $NaH_2PO_4$, 10 mM EDTA, 0.1% Triton and 1 mM 5-bromo-4-chloro-3-indolyl-D-glucuronic acid; pH=7.2) to reveal GUS activity (Silva et al., 2018). After staining, the seedlings were washed in ethanolic series (70%, 80%, 95% and 100%), 1 h for each series at 37 °C to remove chlorophyll. Samples were then photographed using a light microscope (Zeiss AxioScope A1, Jena Germany).

Auxin quantification

The endogenous IAA levels were determined by gas chromatography-tandem mass spectrometry-selecting ion monitoring (Shimadzu model GCMS-QP2010 SE). First, the samples (50-100 mg fresh weight) were extracted and methylated (Rigui et al., 2015). About 0.25 µg of the labeled standard [$^{13}C_6$] IAA (Cambridge Isotopes, UK) was added to each sample as an internal standard. The chromatograph was equipped with a fused-silica capillary column (30 m i.d., 0.25 mm, 0.5-µm-thick internal film) DB-5 MS stationary phase using helium as the carrier gas at a flow rate of 4.5 mL $min^{-1}$ in the following program: 2 min at 100°C, followed by a ramp of 10°C $min^{-1}$ to 140°C, 25°C $min^{-1}$ to 160°C, 35°C $min^{-1}$ to 250°C, 20°C $min^{-1}$ to 270°C and 30°C $min^{-1}$ to 300°C. The injector temperature was 250°C, and the following mass spectrometer operating parameters were used: ionization voltage, 70 eV (electron impact ionization); ion source temperature, 230°C; and interface temperature, 260°C. Ions with mass-to-charge ratios of

130 and 189 (corresponding to endogenous IAA) and 136 and 195 (corresponding to [$^{13}C_6$]IAA) were monitored, and endogenous IAA concentrations were calculated based on extracted chromatograms at mass-to-charge ratios of 130 and 136.

Polar auxin transport assays (PAT)

Hypocotyl sections with 10-mm were excised from MT and *obv* seedlings 2-week-old and incubated in 5 mM phosphate buffer (pH 5.8) containing 1 µM IAA for 2 h at 25°C ± 2°C on a rotary shaker (200 rpm). The segments were placed between receiver blocks (1% [w/v] agar in water) and donor blocks (1% [w/v] agar in 5 mM phosphate buffer [pH 5.8] containing 1 µM IAA and 100 nM [$^3$H] IAA) oriented with their apical ends toward the donor blocks. After 4 h of incubation inside a humid chamber at 25°C ± 2°C, the receiver blocks were removed and stored in a 3 mL scintillation cocktail (Ultima Gold; PerkinElmer). Receiver blocks in the scintillation cocktail were shaken overnight at 100 rpm and 28°C ± 2°C before analysis in a scintillation counter. For the negative control, hypocotyl segments were sandwiched for 30 min between NPA-containing blocks (1% [w/v] agar in water containing 20 µM NPA) an auxin transport inhibitor. $^3$H dpm was converted to fmol of auxin transported (Lewis and Muday, 2009).

Auxin sensitivity assays

For hypocotyl elongation assays, MT and *obv* mutant seeds were grown on half-strength MS medium under photoperiod of 16-h/8-h day/night, light intensity of 45 ± 3 µmol m $^{-2}$ s $^{-1}$ at 25 ± 1 $^{\circ}$C. Hypocotyls from 8-days-old seedlings were excised in sections with 10-mm and preincubated in buffer (10 mM KCl; 1 mM MES-KOH [pH 6] and 1% [w/v] sucrose) for 2 h at 25°C in the dark to deplete endogenous auxin. Then, segments were incubated in the same buffer supplemented with α-naphthalene acetic acid (NAA) in increasing concentrations (0, 0.1, 1, 10, 100 e 1000 µM) for 24 h under agitation, light intensity of 45 ± 3 µmol m $^{-2}$ s $^{-1}$ at 25 ± 1 $^{\circ}$C (Silva et al., 2018). For each genotype, three biological replicates were conducted, each composed of 20 segments. Hypocotyls were photographed and the relative growth rate evaluated by Image Pro-Plus®.

For root regeneration from cotyledon explants, seeds of MT, *obv* mutant and overexpressing lines were germinated *in vitro* in half-strength MS medium under the conditions described above. Cotyledons from 12-days-old seedlings were excised and incubated in MS medium with or without IAA supplementation (0; 0.04; 0.4; 4 and 40 µM). The explants were kept in 25 ± 1 °C under photoperiod of 16-h/8-h and light intensity of 10-20 μmol $m^{-2}$ $s^{-1}$ for 15 days (Wang et al., 2005). The number of explants with visible roots was determined. Four replicates were conducted by genotype, each composed of 12 explants.

Statistical analysis

The experimental design was completely randomized. Data were submitted to analysis of variance (ANOVA) and the means were compared by Tukey test at 5% level of significance ($P \leq 0.05$).

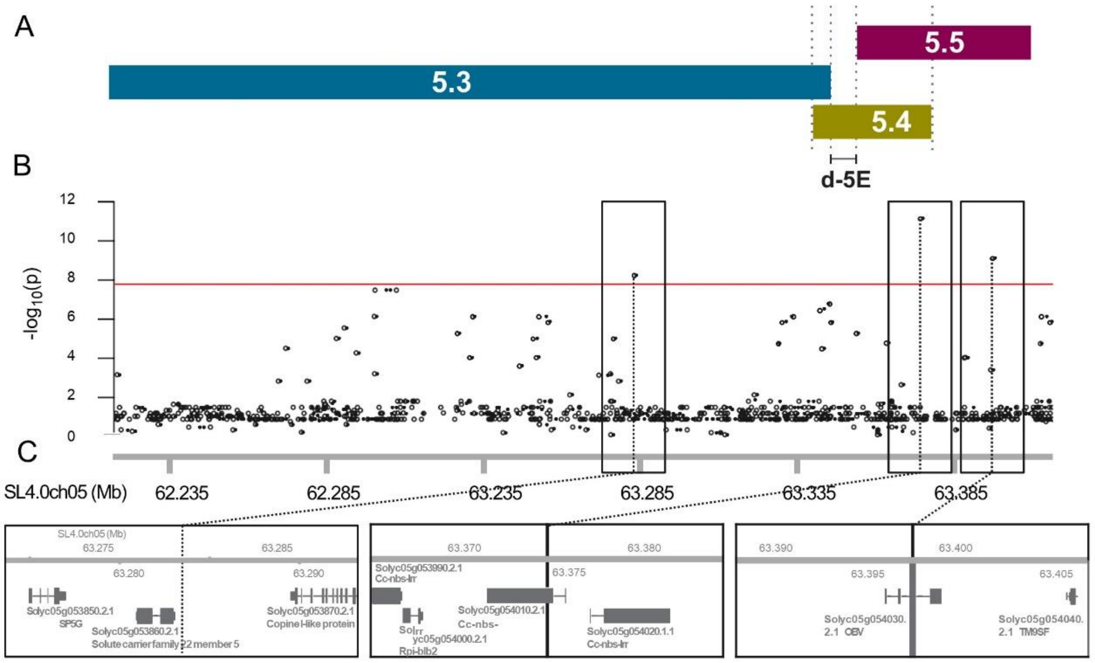


**Figure S1. Mapping and identification of the *OBV* candidate gene.**
(**A**) Schematic depiction of the *Solanum pennellii* segments of the introgression lines in tomato (redrawn from Chitwood et al., 2013) showing the position of bin d-5E (Jones et al., 2007). (**B**) Manhattan plot of the GWAS for *obv* phenotype on d-5E region (63.73 to 64.03 Mb of chromosome 5). X-axis represents the chromosome position and the Y-axis represents the negative log10 of P-values per SNP derived from the association analysis. The red horizontal line represents the Bonferroni-corrected genome-wide significance threshold (P=1.554e-08). (**C**) Gene annotations surrounding SNPs with P-value below the significance threshold are shown in boxes.

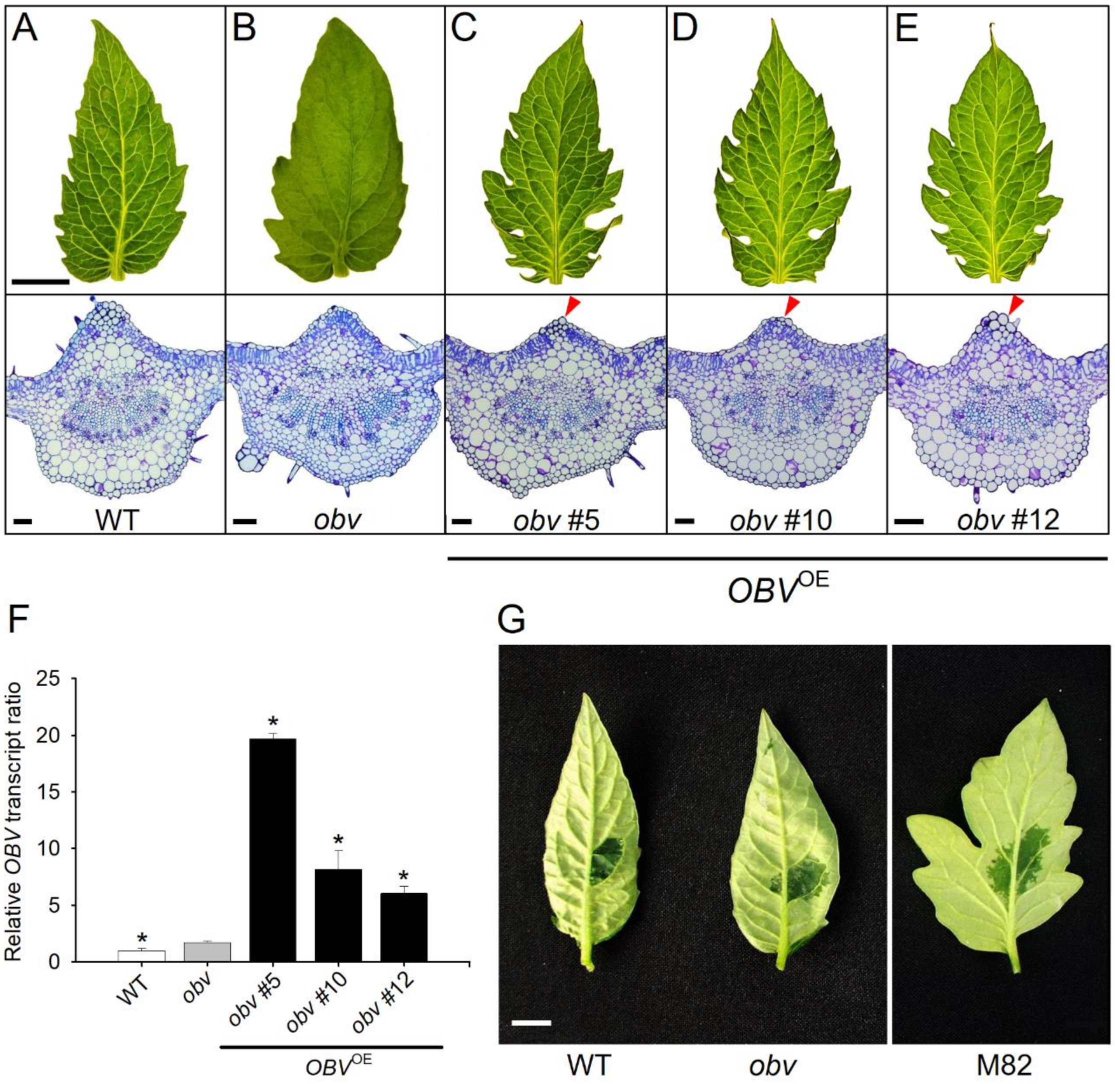


**Figure S2. Complementation of the *obv* mutant with *OBV* functional allele.**
(**A**-**E**) Representative leaflets and cross-sections of Micro-Tom (WT), *obv* mutant and *obv* complemented lines (*OBV*$^{OE}$ #5, #10 and #12) $T_1$ generation. Red arrows show the recovery of BSEs in all *OBV*$^{OE}$ lines. Bars = 1 cm and 100 µm for cross-sections. (**F**) Relative *OBV* mRNA levels in leaves of Micro-Tom (WT), *obv* mutant and three complemented lines (*obv*-*OBV*$^{OE}$ #5, #10 and #12). Values were normalized against the MT (wild-type) sample. Asterisk show significant differences to the *obv* mutant. (**G**) In tomato (WT), BSEs divide the mesophyll into hermetic compartments, creating functionally heterobaric leaves. The compartmentalization of the mesophyll can be illustrated by a quick test, infiltrating water into the leaf blade. In heterobaric leaves, water distribution is limited by barriers imposed by BSEs. Whereas in leaves without the BSEs, homobaric, as in the *obv* mutant and in M82, the artificially infiltrated water spreads freely in the leaf.

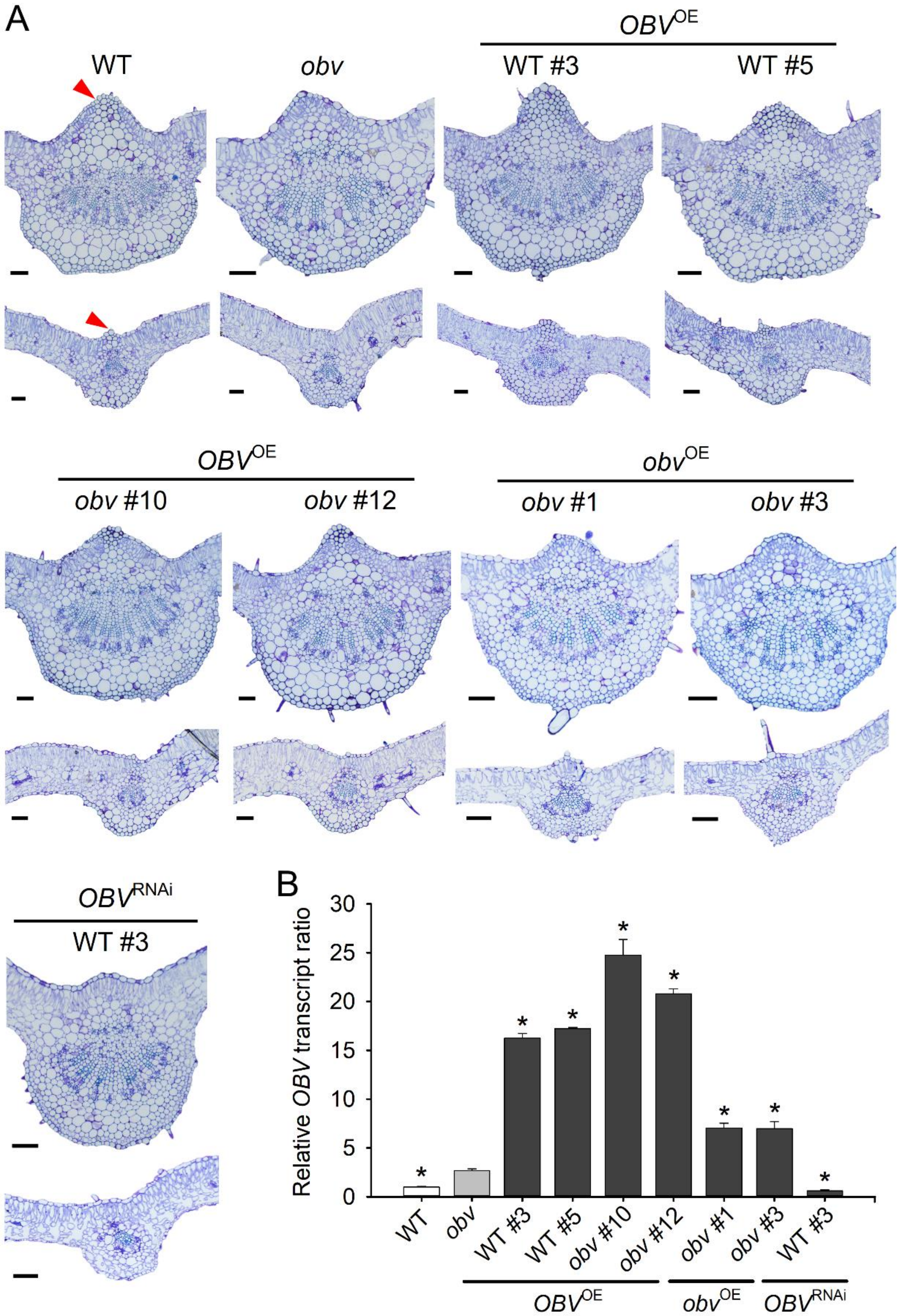


**Figure S3. Complementation of the *obv* mutant and knockdown of *OBV* gene.**

(**A**) Representative cross-sections of primary and secondary veins of Micro-Tom (WT), *obv* mutant, overexpressing lines with a functional allele in a wild-type (WT) (*OBV*$^{OE}$ #3, #5) or mutant (*obv*) (*OBV*$^{OE}$ #10, #12) background; and the mutant allele in an obv mutant background (*obv*$^{OE}$ #1 and #3), and knockdown line in WT background (*OBV*$^{RNAi}$ #3). BSEs (red arrow) are present in all overexpressing lines with *OBV* functional allele, but not in overexpressing lines with *obv* mutant allele. In knockdown line, the reduction in *OBV* expression resulted in complete elimination of BSEs in WT leaves. Bars = 100 µm. (**B**) Relative *OBV* mRNA levels in leaves of WT, *obv* mutant, overexpressing and RNAi lines. Values were normalized against the MT (wild-type) sample. Asterisk show significant differences to the *obv* mutant.

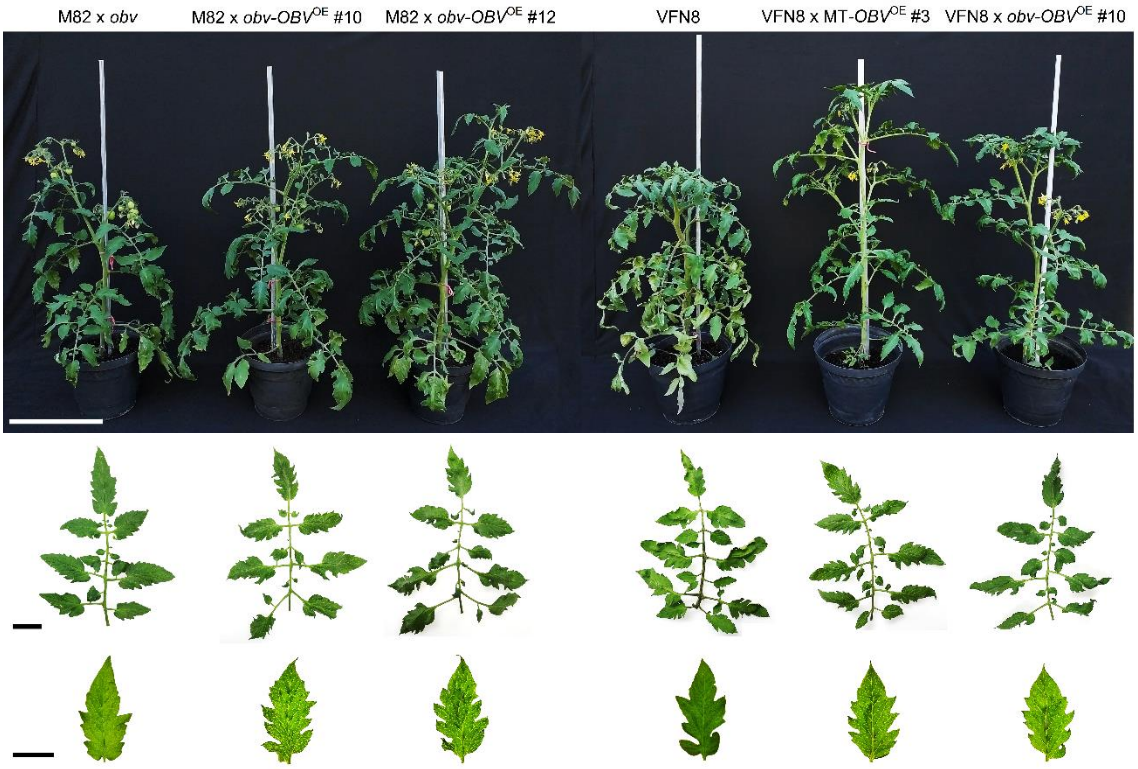


**Figure S4. Morphology of *OBV*-overexpressing (*OBV*$^{OE}$) plants in hybrid M82 × MT and VFN8 × MT backgrounds.** Representative $F_1$ generation plants from crossing of M82 × *obv*, M82 × *obv-OBV*$^{OE}$ #10, M82 × *obv-OBV*$^{OE}$ #12, VFN8, VFN8 × MT-*OBV*$^{OE}$ #3 and VFN8 × *obv-OBV*$^{OE}$ #10. Below are shown leaves and terminal leaflets. Cultivars M82 and VFN8 (both *obv* mutants) has dark veins and the complemented plants with *OBV*$^{OE}$ allele recovered light veins phenotype. Plants were photographed 70 days after germination. Scale bars = 30 cm for plants; = 4 cm for leaves and leaflets.

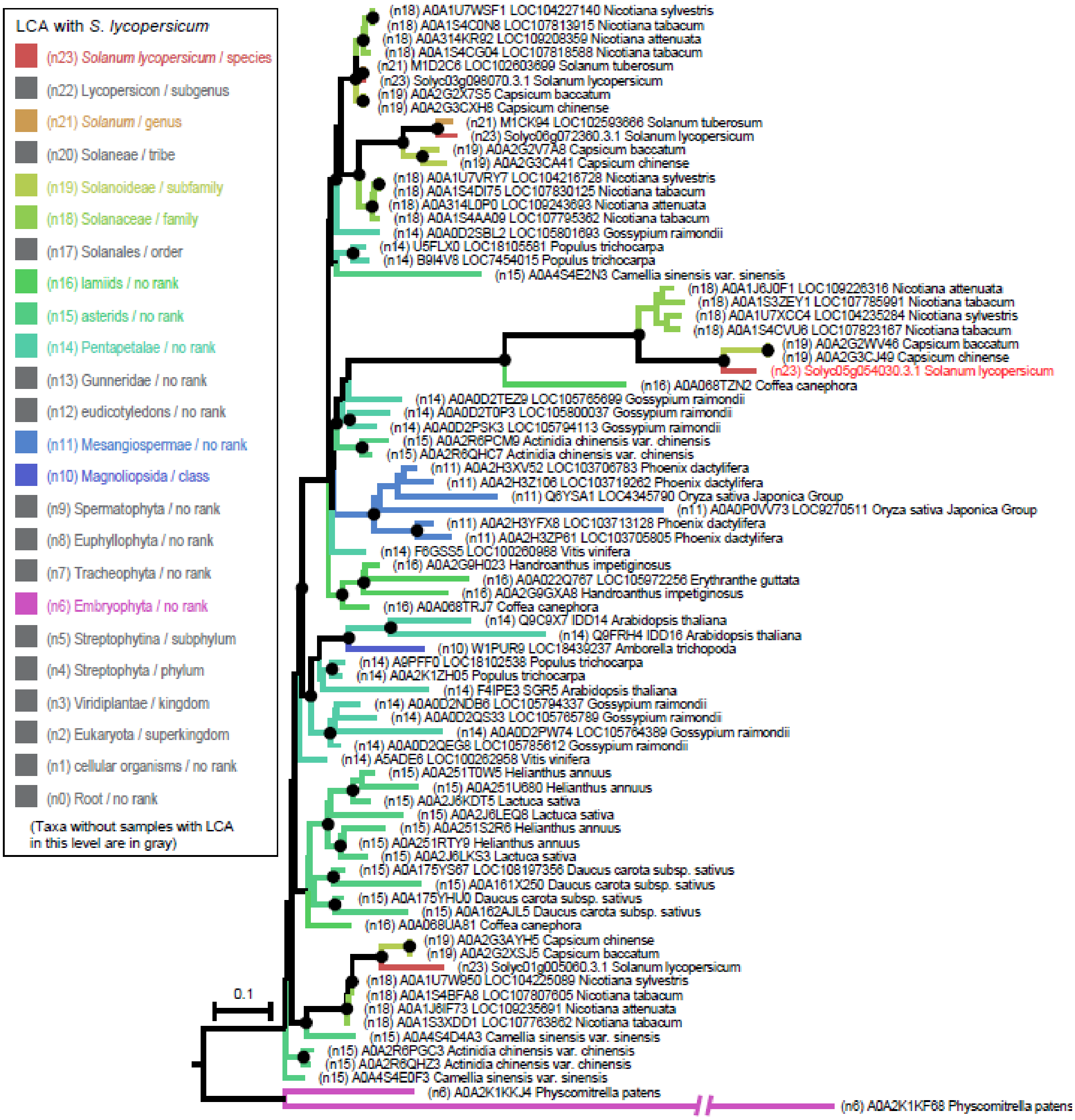


**Figure S5**. **Maximum likelihood tree of OBV family subclade comprising the tomato OBV (in red).** The tree was inferred with PHYML (substitution model: JTT+G+I, bootstrap replicates: 1000). Branches are colored according to the taxonomic relationship between the organism which a sample belongs to with *S. lycopersicum*, which were determined by consulting their Lowest Common Ancestor (LCA) in the NCBI Taxonomy tree. Leaf labels are comprised of (1) LCA level with *S. lycopersicum* (e.g. n11), (2) accession number; (3) gene name when exists and (4) species name.

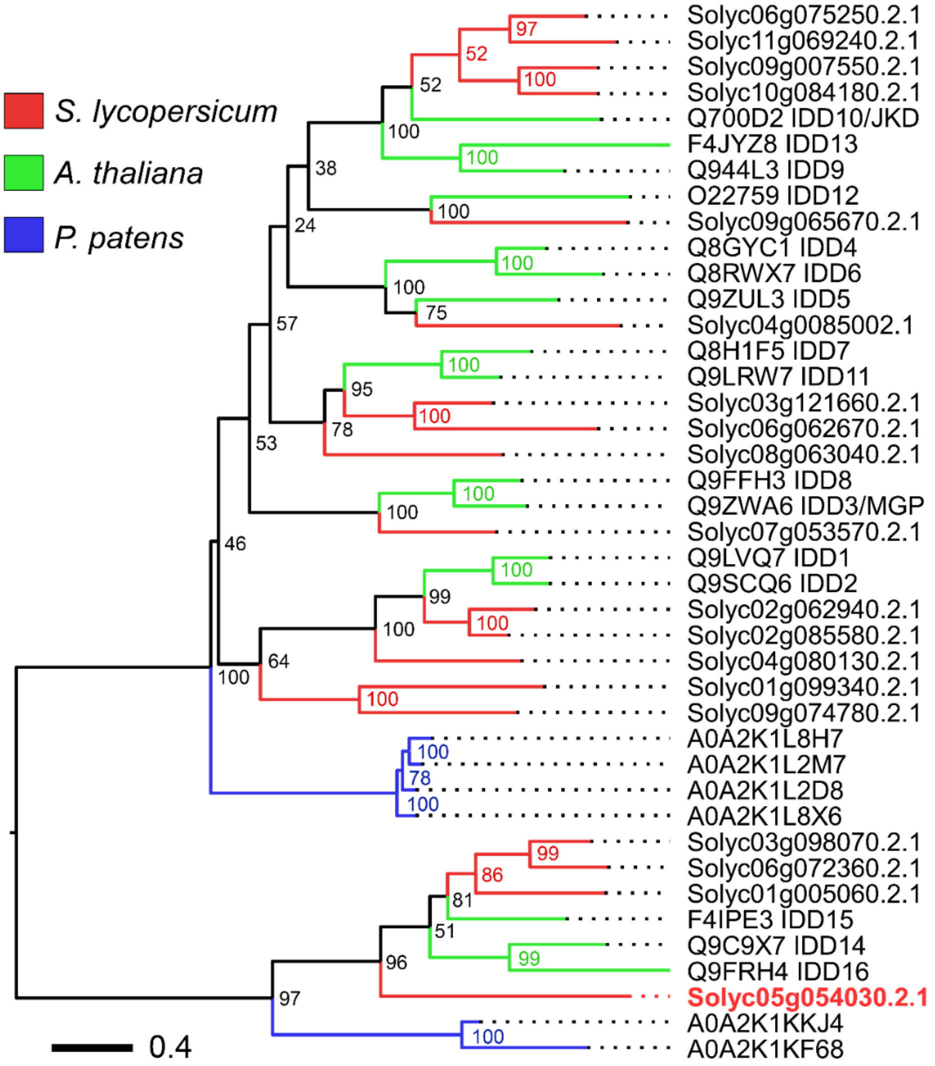


**Figure S6. Phylogenetic reconstruction of the OBV protein family in tomato, Arabidopsis and Physcomitrella patens.** Maximum likelihood tree of the OBV family from *S. lycopersicum*, subclade comprising the tomato OBV (in red) with its paralogs and orthologs in *Arabidopsis thaliana* and the basal angiosperm *Physcomitrella patens*, used as external group. The tree was inferred with PHYML (substitution model: JTT+G+I, bootstrap replicates: 1000). Branch support is shown on each node. Branches are colored according to the organism to which the protein sequence belongs to.

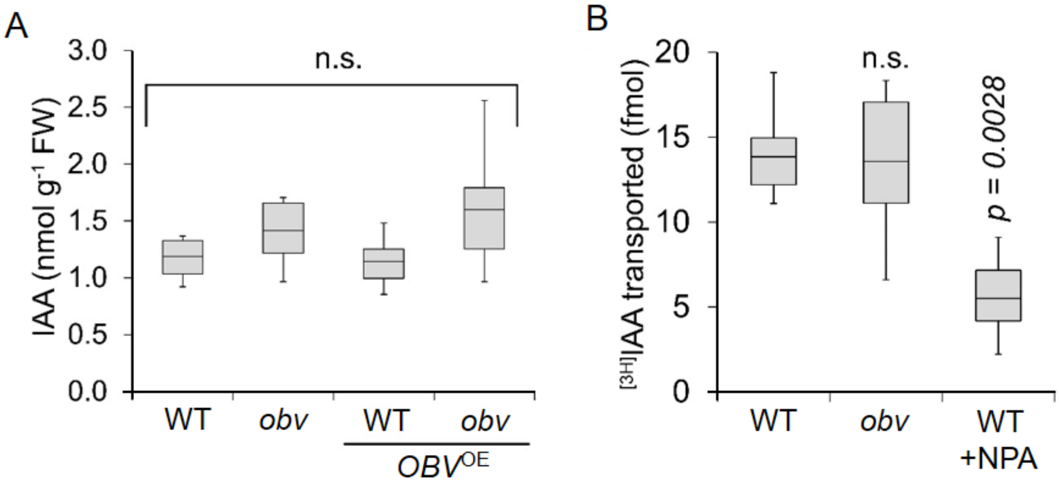


**Figure S7. Influence of *OBV* on leaf free auxin concentration and polar auxin transport in hypocotyl explants.** (**A**) Auxin levels in leaf primordia (≤ 0.5 cm) of MT, *obv* mutant and *OBV*$^{OE}$ lines, homozygous $T_3$. Data represent means of four biological replicates. Each biological replica was composed by a pool of three plants. Different letters indicate significant differences by Tukey's test at 5% probability. FW = Fresh weight. (**B**) [3H]IAA transport in detached hypocotyls of MT and *obv* mutant. The NPA is an auxin polar transport inhibitor and was used as negative control. Data are means (±SE) of at least ten biological replicates.

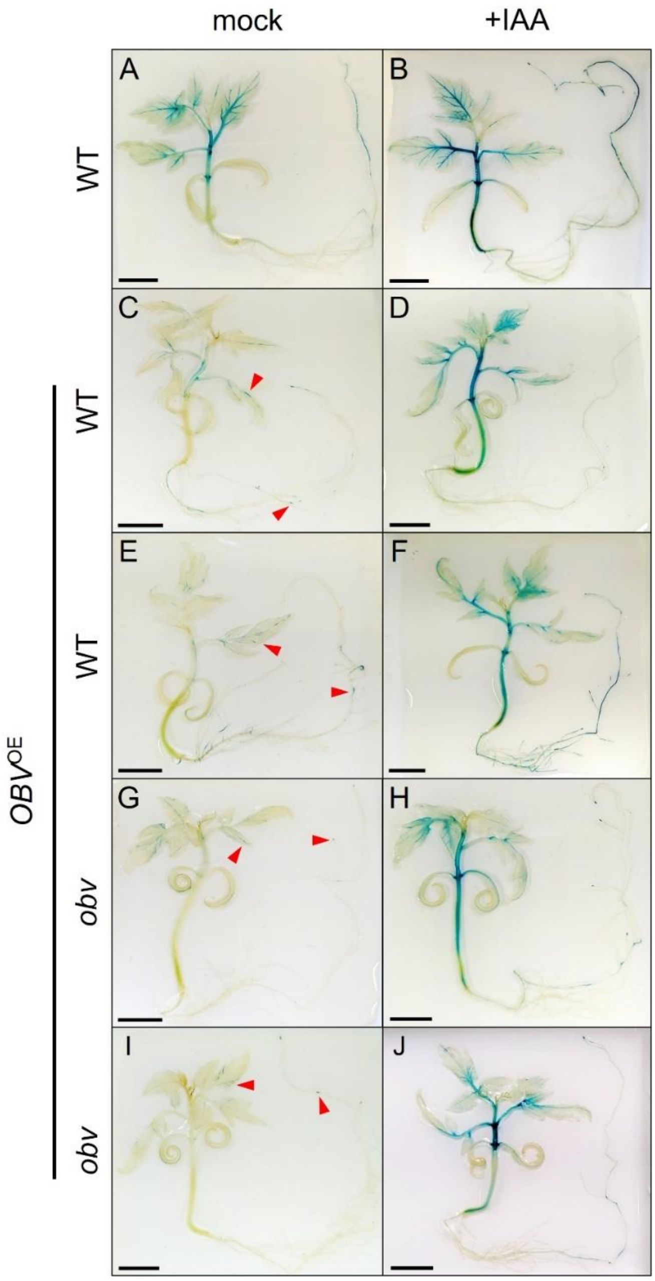


**Figure S8**. **Reduction of auxin sensitivity in *OBV* overexpressing lines revealed by histochemical GUS assays.** Histochemical GUS analysis in MT (MT x *DR5*::*GUS* F1) and overexpression lines (*OBV*$^{OE}$ x *DR5*::*GUS* F1) seedlings at 30 days-old. Expression pattern of *GUS* reporter gene fused to the auxin-inducible *DR5* promoter in wild-type seedlings and *OBV*$^{OE}$ lines (**A**, **C**, **E**, **G**, **I**), that showed a minor sign throughout the seedling. (**B**, **D**, **F**, **H**, **J**) Expression pattern of GUS in MT and *OBV*$^{OE}$ lines treated with exogenous auxin (20 µM IAA). Bars = 1 cm.

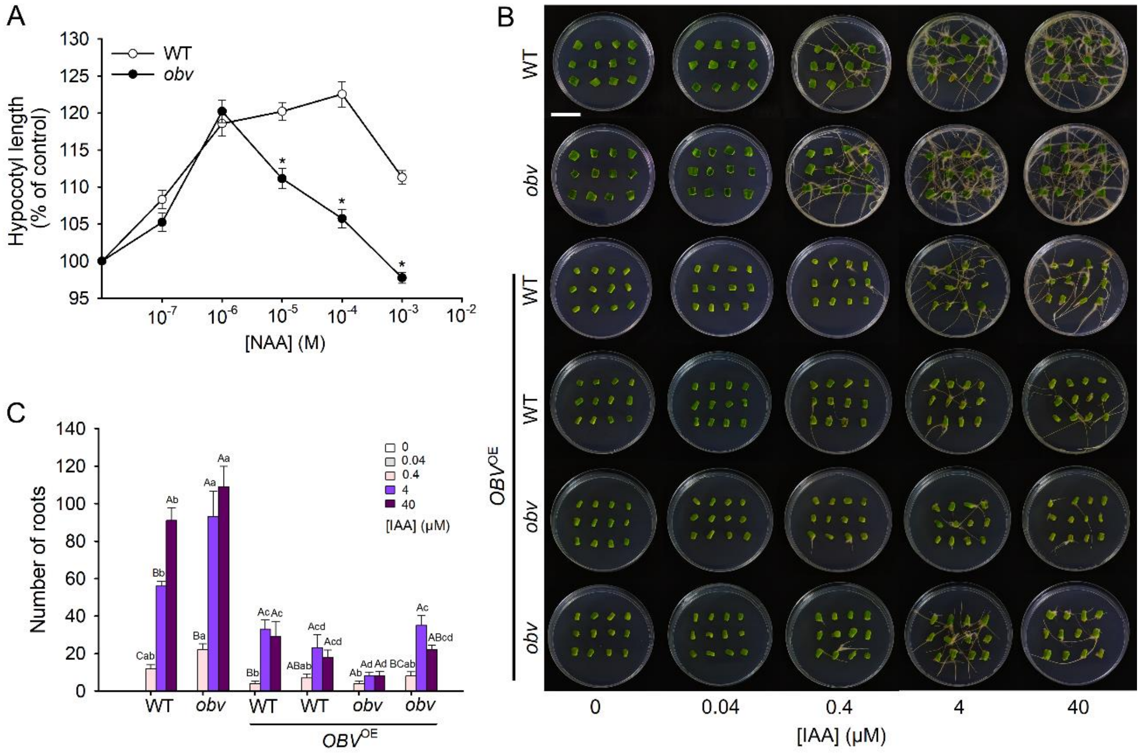


**Figure S9**. **Hypocotyl elongation and *in vitro* rhyzogenesis assays show that OBV alters auxin sensitivity.** (**A**) Auxin dose response in hypocotyl segments. Elongation is given as increase in percentage at final length over the initial length after 24 h incubation in a solution containing the NAA under increasing concentrations. Data are means (±SE) of three biological replicates with ≥ 20 segments for each replicate. Statistically significant differences compared with the MT were determined using Tukey's test: *P<0.05. (**B**) Auxin dose-response assay in 12-day-old cotyledon explants showing a significant reduction in the emission of roots in *OBV*$^{OE}$ lines. (**C**) Total number of emitted roots induced from the concentration of 0.4, 4 and 40 µm IAA, after 15 days of cultivation. Different letters indicate statistically significant differences by Tukey's test, P < 0.05. Uppercase letters compare doses within the same genotype, lowercase letters compare dose effect between different genotypes. Bar = 1cm.

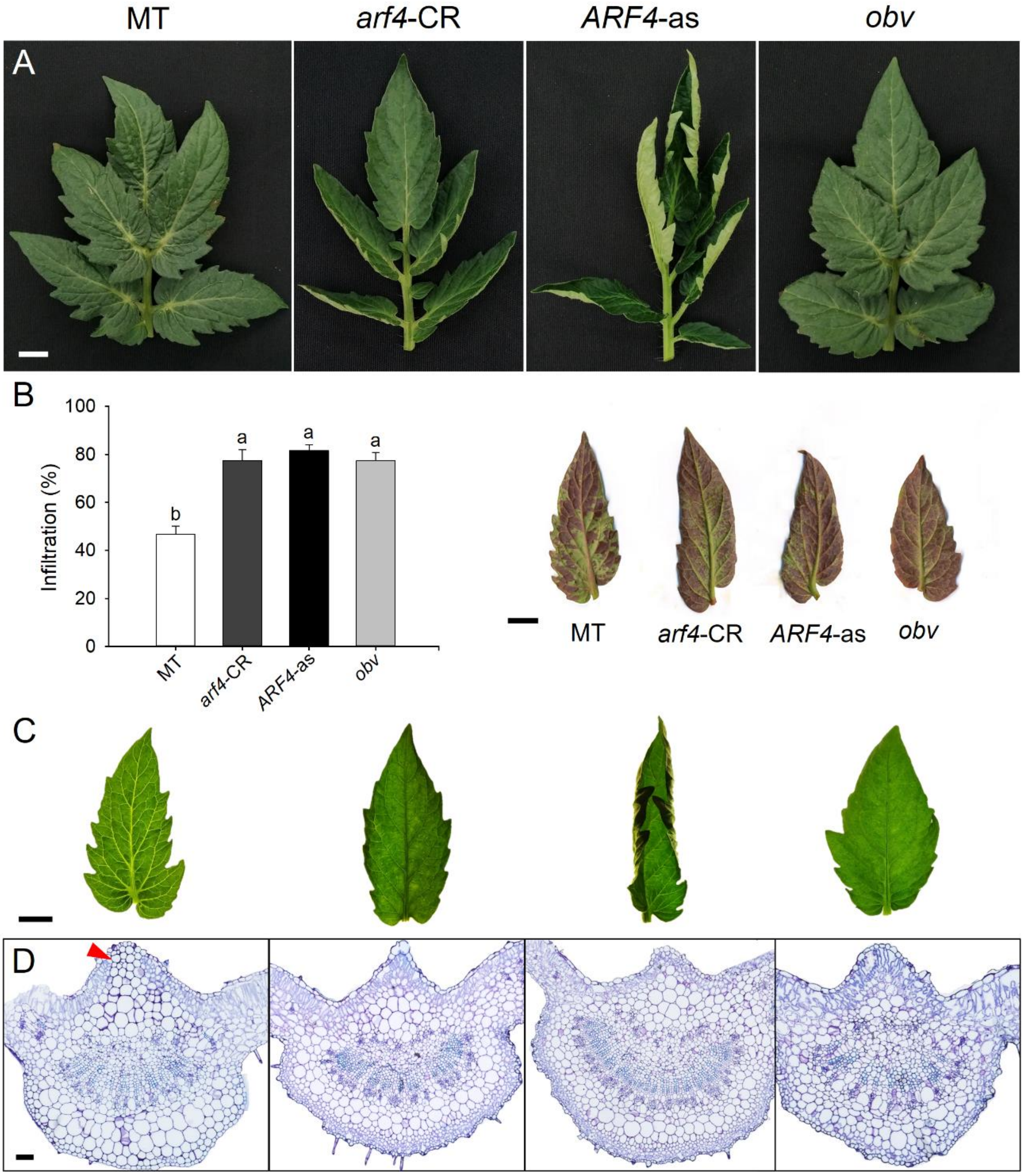


**Figure S10**. **Control of BSEs development by *ARF4* and *OBV*.** (**A**) Representative leaf of MT, *arf4*-CR, *ARF4*-as, and *obv* mutant respectively. (**B**) Infiltration rate shows the percentage of leaf area filled with 1% acid fuchsin dye. The dispersion of the liquid in heterobaric leaves (MT) is hampered by compartmentalization of the mesophyll. In homobaric leaves (*arf4*-CR, *ARF4*-as and *obv*), the liquid is dispersed more easily due to absence of BSEs. The dye infiltration was made with same pressure in detached leaflets. Bars are mean values ± s.e.m. (n=7 leaflets). Different letters indicate significant differences by Tukey's test at 5% probability. (**C**) Details of terminal leaflets under transmitted light evidencing clear or dark vein phenotypes. (**D**) Leaf cross-sections at the midrib showing the presence (MT) and absence (*arf4*-CR, *ARF4*-as and *obv*) of BSEs. Bars = 1 cm and 100 µm for cross-sections.

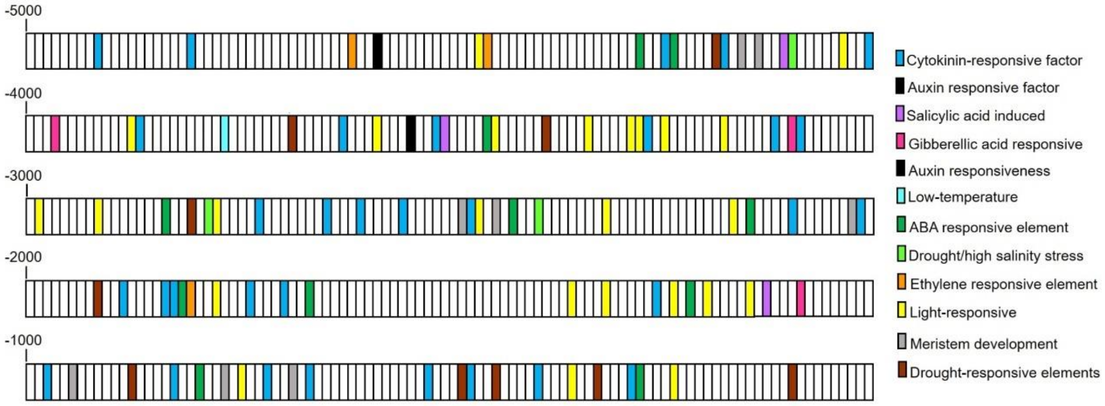


**Figure S11**. ***In silico* analysis of the *OBV* (*Solyc05g054030*) promoter region.** The different motifs are color-coded. See Table S4 for details.

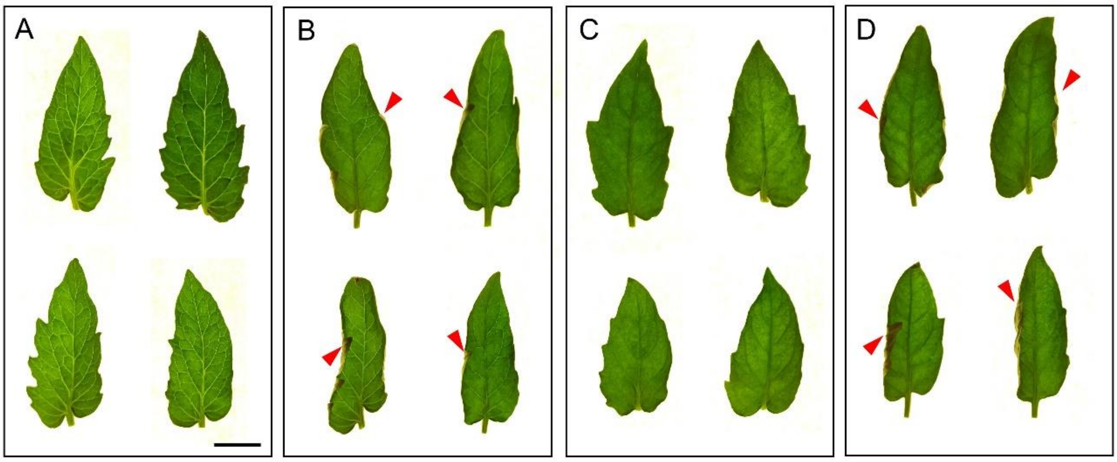


**Figure S12. Phenotypic categories analysed in a dihybrid cross between *arf4* and *obv* mutants.** Representative leaves scored as **(A)** Wild-type, **(B)** *arf4*, **(C)** *obv*, and **(D)** *arf4*, *obv*. The leaf margin inward curling is a characteristic trait of the *arf4* mutant (red arrowheads). The penetrance of the dark vein phenotype is is greater in *obv* than in *arf4*, which allows identification of double mutants based on the presence of leaf curling and entirely dark veins. Scale bar, 1 cm. See Supplementary Table S5 for statistical analysis of the segregation ratios in an $F_2$ population.
